# Tourette disorder features pervasive neuronal and glial transcriptional remodeling in the dorsolateral prefrontal cortex

**DOI:** 10.64898/2026.01.14.699521

**Authors:** Philip J Moos, Caterina Branca, Teresa Musci, Giulia Braccagni, Easton van Luik, Marco Bortolato

**Affiliations:** Department of Pharmacology and Toxicology, College of Pharmacy, University of Utah, Salt Lake City, UT, USA; Department of Cellular and Systems Pharmacology, College of Pharmacy, University of Florida, Gainesville, FL, USA

**Keywords:** Tourette disorder, dorsolateral prefrontal cortex, single-nucleus transcriptomics, transcriptional remodeling, stress-responsive transcription, polygenic risk enrichment

## Abstract

Tourette disorder (TD) is a neurodevelopmental condition with a robust genetic basis, characterized by multiple motor and vocal tics. Tics arise from dysfunction within cortico-striatal-thalamo-cortical circuits, with pathological evidence largely implicating striatal interneuron deficits and microglial activation. Recently, the largest genome-wide association study (GWAS) meta-analysis of TD identified polygenic risk enrichment in Brodmann area 9 (BA9), corresponding to the dorsolateral prefrontal cortex (DLPFC), a region implicated in executive control and tic suppression. However, the molecular landscape of BA9 in TD remains unexplored. Here, we performed single-nucleus RNA sequencing of postmortem BA9 from five males with TD and five matched controls, yielding 72,340 nuclei across neuronal and glial populations. While cell-type proportions were preserved, transcriptional remodeling was pervasive. In particular, biosynthetic and translational programs were upregulated across microglia, interneurons, oligodendrocytes, and superficial- and middle-layer excitatory neurons. Most cell types showed enrichment for glucocorticoid-responsive and immediate early gene modules consistent with stress-associated transcriptional activation. Finally, cross-regional comparison with striatal datasets revealed conservation of microglial and oligodendrocyte programs. These findings point to extensive transcriptional reprogramming in the DLPFC of individuals with TD, characterized by stress-associated activation, most strongly in oligodendrocytes and neurons.

## INTRODUCTION

Tourette disorder (TD) is a neurodevelopmental condition defined by the presence of multiple motor tics and at least one vocal tic persisting for more than one year [1]. Tics are sudden, recurrent movements or vocalizations that are often conceptualized as “unvoluntary” behaviors, insofar as many affected individuals retain a measurable capacity for transient suppression [2, 3]. At the circuit level, tics are widely thought to arise from dysfunction within cortico–striatal–thalamo–cortical (CSTC) loops [4]. Ample postmortem evidence has documented well-defined alterations in the striatum, including reductions in the number of cholinergic and parvalbumin-positive GABAergic interneurons [5–7], as well as microglial activation [7, 8]. These abnormalities have been proposed to generate disinhibitory foci that, when engaged, enable tic expression [9].

Epidemiological and genetic studies further indicate that TD exhibits a marked male predominance [10] and a substantial heritable component. The largest genome-wide association study (GWAS) meta-analysis to date, comprising 9,619 cases and 981,048 controls, identified three genome-wide significant loci mapping to *RBM26, BCL11B*, and *NDFIP2* [11]. Furthermore, this study showed that polygenic risk is enriched in genes preferentially expressed in four CSTC structures, including three striatal areas (putamen, caudate, and nucleus accumbens) and Brodmann area 9 (BA9) of the cortex [11], corresponding to the dorsolateral prefrontal cortex (DLPFC). Several lines of evidence point to a direct implication of this region in the pathophysiology of TD. The DLPFC plays a key role in executive control and the voluntary suppression of tics [12–15]. This region is also characterized by pronounced sensitivity to stress, as catecholamine-mediated weakening of dendritic spines disrupts persistent neuronal firing and reduces regulatory control [16, 17]. This vulnerability is particularly relevant to TD, given that stress is a major modulator of tic severity [18–20].

Taken together, this evidence converges on the DLPFC as a critical yet molecularly uncharacterized node within CSTC circuitry. However, the cellular composition and transcriptional organization of the DLPFC in TD remain largely undefined. To address this gap, we conducted the first single-nucleus RNA sequencing (snRNA-seq) analysis of postmortem BA9 tissue from men with TD and age-matched neurotypical controls. To place these findings in a broader context, we evaluated the extent to which the identified programs intersect with the common variant risk architecture of TD identified in the aforementioned GWAS meta-analysis [11] and assessed cross-regional convergence with striatal transcriptomic data [7].

## RESULTS

The following section summarizes the principal findings. Full statistical details are provided in Supplementary Results.

### Sample collection, processing, and cell-type identification

Postmortem BA9 tissue was collected from 10 male individuals, including 5 with TD and 5 age-matched neurotypical controls (**Table 1**). Nuclei were isolated from frozen tissue by mechanical dissociation, purified through sucrose cushion centrifugation, and processed for 10x Genomics snRNA-seq. One control subject was excluded due to insufficient RNA quality (**Supplementary Fig. 1**), yielding a total of 72,340 high-quality nuclei (29,159 from controls; 43,181 from TD samples).

**Table 1:** Demographic and clinical characteristics of postmortem brain samples included in the study. The cohort comprised ten male subjects, aged 8 to 39 years, with postmortem intervals (PMI) ranging from 6.5 to 30.1 hours. Five subjects had no clinical brain diagnosis and served as controls, whereas five subjects had a confirmed diagnosis of Tourette’s disorder based on clinical records and neuropathological verification. Post-mortem interval (PMI) was calculated as the time between death and brain freezing. All samples were obtained from male donors to minimize confounding effects of sex on gene expression.

| ID | SEX | AGE | PMI | Diagnosis |
| --- | --- | --- | --- | --- |
| NT1 | Male | 19 | 18.58 | No clinical brain diagnosis found |
| NT2 | Male | 33 | 25.67 | No clinical brain diagnosis found |
| NT3 | Male | 24 | 13.1 | No clinical brain diagnosis found |
| NT4 | Male | 18 | 19.83 | No clinical brain diagnosis found |
| NT5 | Male | 35 | 25.67 | No clinical brain diagnosis found |
| TD1 | Male | 39 | 27 | Tourette's disorder (Confirmed Diagnosis) |
| TD2 | Male | 8 | 12.3 | Tourette's disorder (Confirmed Diagnosis) |
| TD3 | Male | 14 | 6.5 | Tourette's disorder (Confirmed Diagnosis) |
| TD4 | Male | 29 | 19 | Tourette's disorder (Confirmed Diagnosis) |
| TD5 | Male | 34 | 30.1 | Tourette's disorder (Confirmed Diagnosis) |

Unsupervised clustering and Azimuth-based annotation identified six major cell types (**Figure 1A-B**): excitatory neurons (23,276 nuclei; 30.8%), interneurons (7,015; 8.3%), astrocytes (9,657; 12.0%), oligodendrocytes (27,321; 36.4%), oligodendrocyte precursor cells (OPCs; 4,392; 6.1%), and microglia (3,093; 4.3%). In contrast to the interneuron depletion documented in the TD striatum [5–7], no significant differences in cell-type proportions were observed between TD and control samples (**Figure 1C**), indicating that cortical pathology in TD operates through transcriptional rather than cytoarchitectural mechanisms.

**Figure 1:**
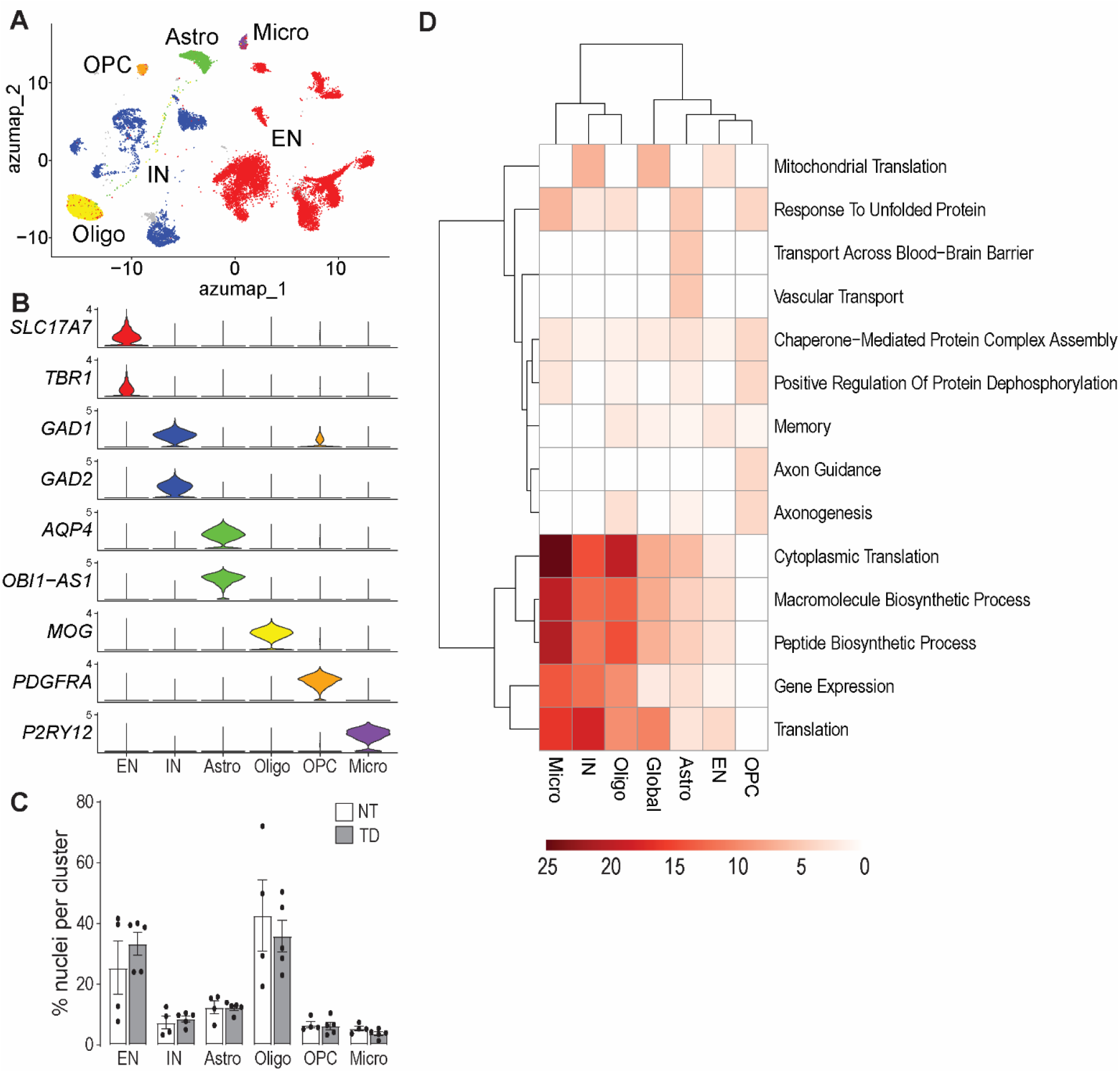
Single-nucleus transcriptomic profiling of BA9 tissues from neurotypical (NT) and Tourette Disorder (TD) patients. (A) Uniform Manifold Approximation and Projection (UMAP) following Azimuth projection on a BA9 reference, annotated by major cell types, including astrocytes, oligodendrocytes, microglia, excitatory neurons, interneurons, and oligodendrocyte progenitor cells (OPC). (B) Violin plots of marker genes used to distinguish the clusters differentiating known cell types. (C) Proportion of nuclei in each cluster, grouped by major cell type, for neurotypical and TD samples. (D) Gene ontology analysis for biological processes highlights the most differential biological processes among these cell clusters. Abbreviations: EN, excitatory neurons; IN, interneurons; Astro, astrocytes; Oligo, oligodendrocytes; OPC, oligodendrocyte precursor cell; Micro, microglia.

### Pervasive transcriptional remodeling with cell-type-specific directionality

Given the limited number of brains, pseudobulk analysis lacked sufficient statistical power; therefore, DEGs were assessed by comparing single-nucleus expression between TD and control brains in the combined dataset. Differential expression analysis revealed extensive transcriptional remodeling across all major cell types, with the direction and magnitude of change varying by population (**Supplementary Data S1-S3**). Interneurons showed the most coherent activation, with 94% of DEGs upregulated (3,227 up, 196 down). Microglia showed a modest upregulatory bias (733 DEGs: 428 up, 305 down). By contrast, excitatory neurons (8,564 DEGs: 3119 up, 5455 down), astrocytes (3,503 DEGs: 537 up, 2,966 down), oligodendrocytes (4,110 DEGs: 815 up, 3,295 down), and OPCs (926 DEGs: 167 up, 759 down) were numerically dominated by downregulation.

Despite this heterogeneity, GO enrichment revealed a unifying functional pattern: biosynthetic and translational programs were consistently upregulated across all major cell types, most prominently in microglia, interneurons, and oligodendrocytes. Astrocytes and microglia additionally showed elevated ribosomal enrichment and RNA- and ubiquitin-binding functions; oligodendrocytes displayed heightened GABA receptor-related signaling; and OPCs exhibited enrichment for unfolded protein response and axon guidance (**Figure 1D** and **Supplementary Fig. 2**). This dissociation between the overall direction of differential expression and the functional significance of the upregulated fraction indicates a generalized shift in the metabolic state of the DLPFC in TD.

### Layer-specific remodeling of excitatory neurons

To determine whether the overall transcriptional pattern was uniform across cortical layers, we examined differential gene expression across excitatory neuron subtypes. These cells were resolved into six layer-specific clusters (**Figure 2A-B**): L2-3 Superficial Intratelencephalic (IT), L3-4 Middle IT, L5-6 Deep IT, L5/6 Near-Projecting (NP), L6 Corticothalamic (CT), and L6b neurons. No significant differences in proportions were found between groups (**Figure 2C**).

**Figure 2:**
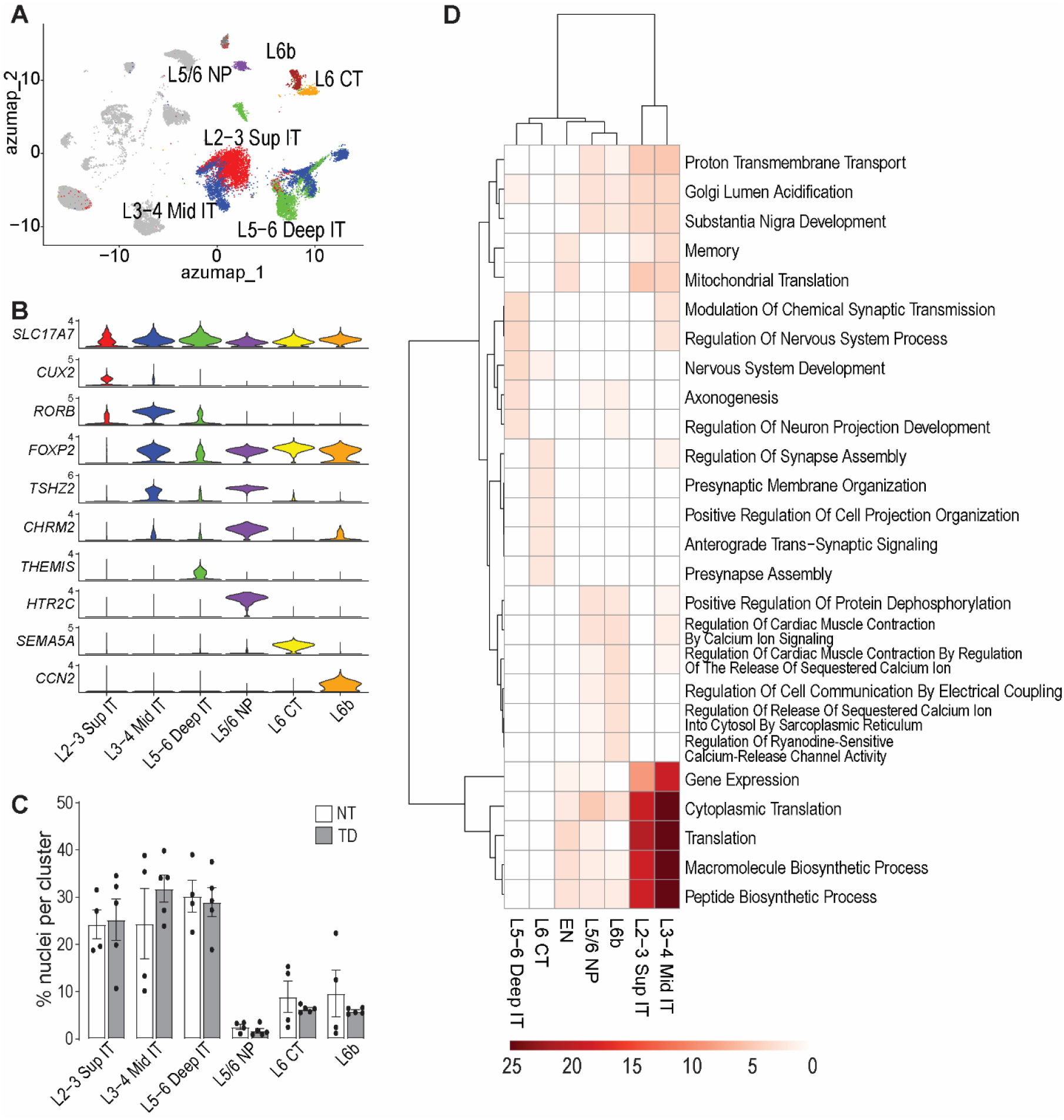
Cluster analysis of excitatory neurons from neurotypical (NT) and Tourette Disorder (TD) patients. (A) Uniform Manifold Approximation and Projection (UMAP) visualization after Azimuth projection and annotated by major cell types. (B) Violin plots showing the expression of representative layer-specific marker genes used to define the clusters. (C) Proportion of nuclei in each cluster, grouped by major cell type, for NT and TD samples. (D) Gene ontology analysis for biological processes highlights the most differential biological processes among these cell clusters. Abbreviations: EN, excitatory neurons; L2-3 Sup IT, Layer 2-3 Superficial Intratelencephalic; L3-4 Mid IT, Layer 3-4 Middle Intratelencephalic; L5-6 Deep IT, Layer 5-6 Deep Intratelencephalic; L5/6 NP, Layer 5/6 Near-Projecting neurons; L6 CT, Layer 6 Corticothalamic; and L6b, Layer 6b.

A pronounced laminar divergence emerged (**Supplementary Data S4-S6**). Superficial (L2-3) and middle (L3-4) layers were dominated by downregulated genes, whereas deep layers showed a predominance of upregulated genes. Remarkably, functional enrichment inverted this pattern (**Figure 2D** and **Supplementary Fig. 3**): biosynthetic and translational pathways were upregulated in superficial and middle layers despite their overall transcriptional suppression, whereas deep layers showed downregulation of these same pathways despite their overall transcriptional activation. Deep-layer neurons instead engaged developmental and synaptic remodeling programs. This laminar dissociation maps onto the functional architecture of the DLPFC, superficial IT neurons mediate corticocortical communication, middle-layer neurons occupy the primary thalamocortical recipient zone processing feedback from the basal ganglia, and deep-layer neurons maintain corticostriatal and corticothalamic projections [21], suggesting that TD remodeling is not a uniform transcriptional shift but a layer-specific reorganization tied to distinct circuit roles. Consistent with this interpretation, pseudotime analysis revealed that TD excitatory neurons occupied earlier positions along inferred maturation trajectories (**Supplementary Fig. 4** and **Supplementary Data S7**), indicating that the layer-specific transcriptional changes reflect a broader pattern of disrupted neuronal maturation, one that aligns with the childhood onset and developmental trajectory characteristic of TD.

### Widespread transcriptional activation in interneurons

All four interneuron subtypes, distinguished by canonical markers (**Figure 3A-C**), showed near-exclusive upregulation of DEGs (**Supplementary Data S8-S10**): VIP+ (515 up, 51 down), LAMP5+ (263 up, 27 down), PV+ (1,318 up, 97 down), and SST+ (496 up, 29 down). This activation contrasts sharply with the mixed directionality observed in excitatory neurons and glial populations.

**Figure 3:**
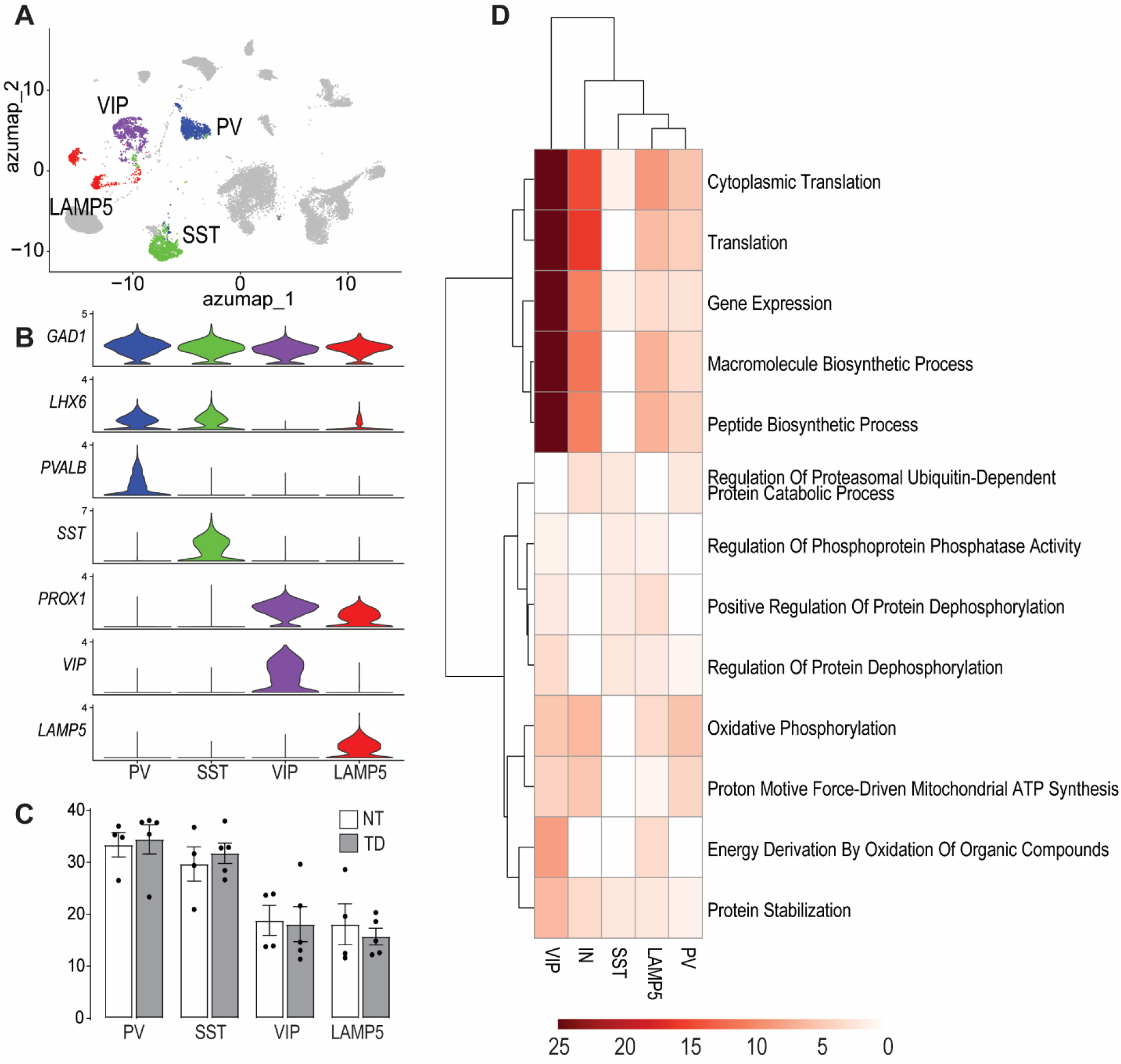
Cluster analysis of interneurons from neurotypical (NT) and Tourette Disorder (TD) patients. (A) Uniform Manifold Approximation and Projection (UMAP) visualization after Azimuth projection and annotated by major cell types. (B) Violin plots showing the expression of representative specific marker genes used to define the clusters. (C) Proportion of nuclei in each cluster, grouped by major cell type, for NT and TD samples. (D) Gene ontology analysis for biological processes highlights the most differential biological processes among these cell clusters. Abbreviations: SST, somatostatin-positive; PV, parvalbumin-positive.

GO analyses revealed shared and subtype-specific signatures (**Figure 3D** and **Supplementary Fig. 5**). VIP+ and LAMP5+ interneurons showed the strongest enrichment for biosynthetic and translational programs. VIP+ activation is particularly notable given the role of these cells in disinhibitory microcircuits, where they suppress SST+ and PV+ interneurons to release pyramidal neurons from inhibition. PV+ interneurons combined translational activation with oxidative phosphorylation and mitochondrial enrichment, while SST+ cells were enriched for proteostatic programs. Trajectory inference further placed TD interneurons at earlier positions along pseudotime-ordered maturation paths, indicating that the observed transcriptional changes may reflect a disruption in developmental timing rather than a stable mature state (**Supplementary Fig. 6** and **Supplementary Data S11**).

### Stress-responsive transcriptional signatures

The coordinated biosynthetic activation across all cell types, despite cell-type-specific differences in overall directionality, resembles the prefrontal transcriptional response to sustained stress. Because the DLPFC is a well-established target of glucocorticoid-mediated remodeling [22], and because stress is a recognized modulator of tic severity [18], we examined whether TD-associated signatures overlapped with known stress-responsive programs, querying two curated gene sets representing glucocorticoid-responsive [23] and immediate early gene (IEG) [24] modules.

Comparison with glucocorticoid-responsive modules derived from dexamethasone-exposed iPSC neurons [23] revealed widespread enrichment across all major DLPFC cell populations in TD (**Figure 4A** and **Supplementary Data S12**). *CRH*, encoding corticotropin-releasing hormone, was markedly upregulated in interneurons (log2FC = 0.91, padj = 1.6 × 10^−11^), with the strongest induction in PV+ (log2FC = 1.35) and VIP+ cells (log2FC = 1.11), where over half of TD nuclei expressed detectable transcript compared with approximately 30% in controls. Leveraging the same reference dataset [23], which also characterizes DLPFC transcriptional signatures in MDD and PTSD, we found substantial cross-disorder overlap with TD-associated DEGs across multiple cell types (most prominently in excitatory neurons and interneurons) with UMAP projections confirming that shared transcriptional responses localized to the same clusters upregulated in TD, suggesting convergent stress-mediated prefrontal dysfunction rather than nonspecific consequences of psychopathology (**Supplementary Fig. 7-8, Supplementary Table 1**, and **Supplementary Data S13**).

**Figure 4.**
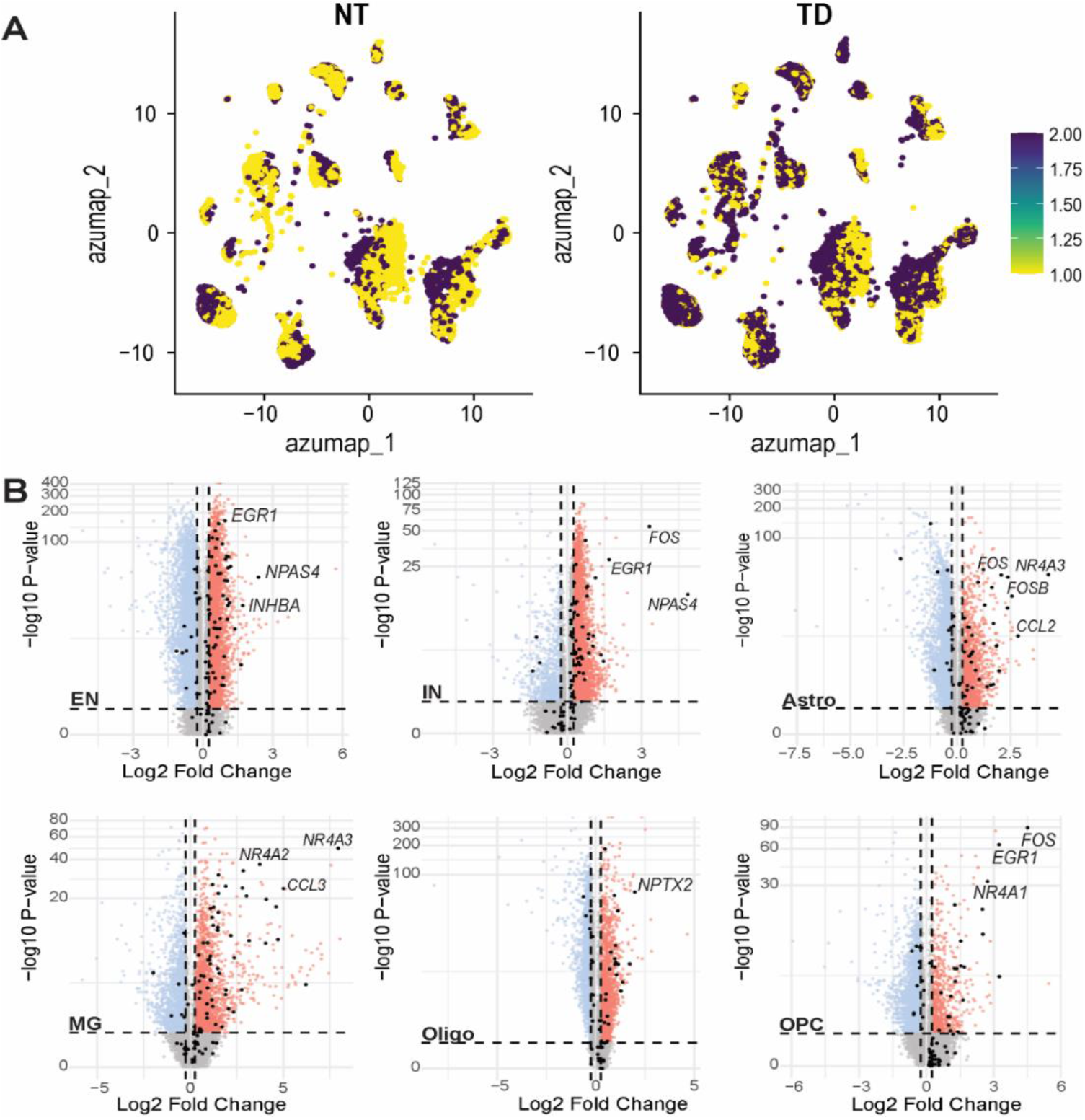
Cell-type-specific IEG activation signatures in Tourette disorder. (A) Dexamethasone-responsive gene module scores for neurotypical (NT) and Tourette disorder (TD) conditions. Colors represent module expression levels from low (yellow) to high (dark purple) based on dexamethasone-induced gene signatures derived from [23]. Each dot represents an individual cell positioned by transcriptional similarity. (B) Volcano plots showing differential gene expression analysis for six major brain cell types: excitatory neurons (EN), inhibitory neurons (IN), astrocytes (Astro), microglia (MG), oligodendrocytes (Oligo), and oligodendrocyte precursor cells (OPC). X-axis displays log_2_ fold change (TD vs. NT), Y-axis shows −log_10_ P-value. Red dots denote significantly upregulated DEGs, blue dots denote significantly downregulated DEGs, and gray dots represent genes that do not meet the significance or fold-change thresholds (|log_2_FC| < 0.25 and P ≥ 0.05). Black dots indicate immediate early genes (IEGs). Key IEGs are labeled, including *EGR1, FOS, FOSB, NR4A3, NR4A2, CCL2, NPTX2*, and *NR4A1*. Dashed horizontal line represents P = 0.05 significance threshold; dashed vertical lines indicate ±0.25 log_2_ fold change thresholds. IEG enrichment analysis conducted using curated reference gene set from [24].

Using a curated IEG reference list [24], we identified strong enrichment across multiple TD cell populations, most prominently in microglia, followed by astrocytes and OPCs (**Supplementary Table 2** and **Supplementary Data S14**). A consistent pattern of coordinated upregulation emerged for *NR4A1, NR4A2, NR4A3, FOS, FOSB*, and *EGR1* across both neuronal and glial populations (**Figure 4B**). Because IEG expression can be influenced by perimortem conditions, we assessed potential confounds; PMI did not differ between groups, showed no significant correlation with IEG expression at either the donor or cell-type level, and single-nucleus PMI correlations were predominantly negative, opposite to the direction expected from agonal artifact.

### Convergence with the genetic architecture of TD

We next evaluated whether these alterations intersect with the common-variant genetic architecture of TD, as predicted by the GWAS enrichment of polygenic risk in BA9-expressed genes [11]. Notably, all three genes reaching significance in the GWAS gene-based analyses were differentially expressed in our dataset: *BCL11B* (MAGMA z = 5.39, the strongest gene-level signal in the entire TD GWAS) and *NDFIP2* were upregulated, while *RBM26* was downregulated.

Overlap with eQTL- and sQTL-linked genes revealed a striking dissociation (**Supplementary Figure 9A–B** and **Supplementary Data S15**): DEGs were uniformly depleted among eQTL-linked genes (upregulated: ORs 0.41-0.75; downregulated: ORs 0.65-0.90), yet consistently enriched among sQTL-linked genes (upregulated: ORs 1.07-1.96; downregulated: ORs 1.40-1.87), suggesting that TD risk variants preferentially influence splicing rather than expression levels of transcriptionally altered genes.

MAGMA analyses using conventional (c-MAGMA), chromatin interaction-informed (h-MAGMA), and eQTL-informed (e-MAGMA) approaches (**Supplementary Data S15**) revealed enrichment exclusively among upregulated DEGs under c-MAGMA and h-MAGMA, concentrated in OPCs, microglia, oligodendrocytes, and inhibitory neurons (ORs 1.27-1.97). e-MAGMA replicated this signal but additionally revealed broad enrichment among downregulated DEGs across most cell types (ORs 1.24-1.45), likely reflecting broadly expressed regulatory variants detectable only through eQTL-based mapping.

BrainSMR analyses revealed a further asymmetry (**Supplementary Figure 9A–B** and **Supplementary Data S16**). For upregulated DEGs, enrichment was concentrated in the oligodendrocyte lineage (with GWAS-implicated oligodendrocyte genes: OR = 2.34, P = 5.5 × 10^−8^; OPC/COP genes: OR = 2.47, P = 4.6 × 10^−4^). For downregulated DEGs, broad enrichment was observed across nearly all cell types (microglia: OR = 8.40; OPCs: OR = 6.01; oligodendrocytes: OR = 3.37; excitatory neurons: OR = 2.57; astrocytes: OR = 2.89; all P < 10^−11^), except inhibitory neurons (OR = 2.12, P = 0.072), consistent with a genetically constrained transcriptional landscape.

Together, these analyses reveal structured intersection between DLPFC transcriptomic signatures and TD genetic risk, organized by three principles: (1) splicing over expression regulation; (2) cell-type-specific GWAS enrichment for upregulated DEGs in glial and inhibitory neuron populations versus broad, conditioning-dependent enrichment for downregulated DEGs; and (3) an asymmetry whereby upregulated genes converge with genetic risk selectively in oligodendrocytes while downregulated genes show convergence across nearly all cell types. These findings suggest that TD risk variants shape the DLPFC transcriptional landscape through post-transcriptional mechanisms, with glial and inhibitory neuron activation programs particularly proximal to genetically mediated disease pathways.

### Cross-regional comparison with striatal transcriptomics

Finally, to determine whether the transcriptional changes in the DLPFC reflect region-specific or shared pathological processes across CSTC circuitry, we compared our BA9 DEGs with published striatal snRNA-seq data from TD [7].

Glial populations showed the strongest cross-regional conservation (**Supplementary Fig. 10** and **Supplementary Data S17-S20**). Microglia exhibited the highest convergence (OR = 20.2), with shared signatures spanning immune activation, lipid metabolism, and endolysosomal trafficking. Oligodendrocytes converged on calcium signaling, vesicular trafficking, and mitochondrial stress (76 shared GO terms). Astrocytes shared neurovascular and lipid metabolism programs. OPCs exhibited conserved synaptic interface organization and innate immune activation. In contrast, neuronal changes were largely region-specific, with BA9 and striatal interneurons converging primarily on downregulated synaptic structure programs (synaptic vesicle cycle OR = 51.0, presynapse assembly OR = 25.6). These patterns suggest that shared glial activation constitutes the biological substrate linking cortical and subcortical pathology in TD, whereas neuronal alterations are specific to each CSTC node.

## DISCUSSION

This study provides the first single-nucleus transcriptomic characterization of the DLPFC in TD. Profiling over 72,000 nuclei, we delineated a cortical molecular landscape defined by four principal features: (i) preservation of cellular composition alongside widespread transcriptional remodeling; (ii) laminar- and cell-type-specific reorganization of neuronal gene expression; (iii) coordinated activation of biosynthetic, metabolic, and stress-responsive programs across neuronal and glial compartments; and (iv) structured convergence with the common-variant genetic architecture of TD. Collectively, these findings indicate that cortical pathology in TD is characterized by sustained transcriptional engagement rather than overt cell loss.

All major cortical cell classes were represented in comparable proportions between TD and control samples, in contrast to the striatal interneuron depletion documented in histopathological [5, 6] and transcriptomic [7, 8] studies. This dissociation suggests that interneuron loss represents a region-specific vulnerability of subcortical nodes, whereas the DLPFC exhibits a distinct pathological mode defined by transcriptional remodeling within an intact cellular architecture. Cortical and striatal components of CSTC circuitry therefore contribute to TD through mechanistically distinct processes that remain functionally coupled at the circuit level.

Within excitatory populations, laminar organization revealed a bidirectional pattern. DEGs were downregulated in superficial (L2–3) and middle-layer (L3–4) IT neurons and upregulated in deep-layer projection neurons. Functional enrichment analyses showed the opposite trend, with GO terms enriched among upregulated genes in superficial and middle layers and among downregulated genes in deep layers. These three laminar compartments correspond to distinct positions within CSTC circuitry [25]. L2-3 superficial IT neurons mediate corticocortical communication, including regulatory projections to premotor areas; L3-4 middle IT neurons occupy the primary thalamocortical recipient zone and process feedback from the basal ganglia via the mediodorsal thalamus; and L5-6 deep IT neurons maintain corticostriatal and corticothalamic projections [21]. Biosynthetic activation in superficial and middle layers may reflect the sustained demands of tic suppression and chronic engagement of the thalamocortical input stage, consistent with a tonically overactive CSTC loop [4]. By contrast, transcriptional activation of developmental and synaptic programs in deep layers may correspond to enhanced excitatory output to subcortical targets implicated in tic generation. This laminar dissociation provides a plausible molecular substrate for the coexistence of executive dysfunction and excessive motor output in TD, with middle-layer neurons serving as a potential indicator of loop-level circuit demand.

Interneuron populations exhibited coherent transcriptional activation across all major subclasses. VIP+ interneurons displayed the strongest enrichment for biosynthetic and translational programs, consistent with their established role in disinhibitory microcircuits.

The activation of VIP+ interneurons also provided a potential mechanistic bridge between disinhibitory and stress-responsive processes. *CRH* was markedly upregulated in VIP+ cells, consistent with prior evidence that these neurons are a key cortical source of this neuropeptide [26, 27]. VIP and CRF act through distinct receptors (VPAC1 and CRF1) to modulate pyramidal cell excitability via cAMP/PKA signaling [28]. Thus, activated VIP+ interneurons may both disinhibit pyramidal neurons (via suppression of SST+ and PV+ interneurons) and directly enhance their excitability through CRF signaling. Given the preferential localization of VIP+ interneurons to superficial layers [29], enhanced *CRH* signaling may contribute to the selective biosynthetic activation observed in superficial-layer excitatory neurons. Concurrent *CRH* upregulation in PV+ interneurons, accompanied by increased *CRHBP* expression, further supports engagement of stress-responsive neuropeptide signaling across inhibitory populations, with local regulation of ligand availability.

Pseudotime analyses indicated that TD neurons occupied earlier positions along inferred maturation trajectories, particularly within excitatory populations. Whether this reflects primary developmental alterations or secondary adaptations to chronic circuit engagement remains unresolved; however, it raises the possibility that altered maturation dynamics contribute to the developmental course of TD.

Across neuronal and glial populations, TD samples exhibited coordinated upregulation of protein synthesis, ribosomal function, and mitochondrial metabolism, consistent with sustained energetic demand. This pervasive upregulation supports a model of persistent circuit engagement rather than static dysfunction. Notably, overlap with MDD and PTSD transcriptional signatures, including enrichment for glucocorticoid-responsive genes and IEG modules (*FOS, EGR1, NR4A* family), likely reflects common downstream consequences of sustained prefrontal activation in conditions imposing chronic demands on executive control. These molecular signatures provide a plausible substrate for the well-established sensitivity of tic expression to stress [18–20].

Integration with the genetic risk architecture further supported the relevance of these transcriptional programs. All three GWAS-identified genes (BCL11B, NDFIP2, RBM26) were differentially expressed, with *BCL11B* and *NDFIP2* specifically upregulated in TD. Differential expressed genes were enriched among sQTL-but not eQTL-linked genes, suggesting that TD risk preferentially influences splicing. Upregulated genes showed cell-type-specific MAGMA and brainSMR enrichment concentrated in oligodendrocytes, OPCs, microglia, and inhibitory neurons.

Cross-regional analyses demonstrated that glial, rather than neuronal, transcriptional programs were preferentially conserved across CSTC nodes. Microglia exhibited the strongest convergence, with shared signatures involving immune activation, lipid metabolism, and endolysosomal trafficking. Oligodendrocytes converged on calcium signaling, vesicular trafficking, and mitochondrial stress. This breadth of glial convergence suggests that metabolic and inflammatory processes constitute a circuit-wide substrate linking cortical and subcortical pathology, whereas neuronal alterations are comparatively region-specific. Cross-regional interneuron convergence was limited to downregulated synaptic structure programs, suggesting a shared state in which vesicular processing is preserved while inhibitory synaptic architecture is progressively compromised.

The convergence of transcriptomic, genetic, and cross-regional findings supports a provisional model, which we present as a framework for guiding future investigation rather than as a definitive causal account.

Whether this transcriptional state reflects a primary genetic predisposition, a compensatory response, or both, cannot be determined from postmortem data. GWAS convergence favors a primary component, whereas glucocorticoid and IEG enrichment suggests experience-dependent amplification. In this framework, the DLPFC contributes to TD through two complementary mechanisms: enhanced corticostriatal drive from deep layers that may facilitate tic generation, and chronic metabolic strain on superficial-layer regulatory circuits that may degrade the precision of tic suppression, with stress exacerbating both through *CRH*-mediated pathways.

Postmortem analyses do not permit causal inference, and the model remains hypothetical until tested prospectively. snRNA-seq captures nuclear transcripts and may not fully reflect cytoplasmic RNA or protein levels. Thus, the relationship between the transcriptional signatures documented here and actual protein output or circuit-level activity remains to be established. Medication histories were unavailable, and subclinical comorbidities, particularly ADHD and OCD, could contribute to the observed signatures. The cohort included only male donors, and the modest sample size (n = 5 TD, n = 4 controls) limits statistical power and precludes examination of sex differences. Circuit-level interpretations, particularly regarding VIP+/*CRH*-mediated disinhibition and corticostriatal overdrive, are inferred from transcriptional data and require electrophysiological and pharmacological validation.

Testable predictions arising from the model include (i) targeted manipulation of VIP+ interneuron activity and CRF1 signaling in TD animal models, (ii) correlation of TD polygenic risk scores with prefrontal activation measures in imaging cohorts, (iii) assessment of CSTC tract myelination as a function of genetic load, and (iv) examination of whether chronic corticostriatal overdrive produces selective striatal interneuron vulnerability. Replication in larger and more diverse cohorts, including female donors and individuals across the tic severity spectrum, will be essential to assess generalizability.

Despite these limitations, this study establishes a cellular and molecular framework for understanding DLPFC involvement in TD. The cortex is not a passive node in CSTC circuitry but an actively remodeled structure, genetically predisposed toward transcriptional hyperactivation and further shaped by the chronic demands of the disorder.

## METHODS AND MATERIALS

The following descriptions provide a summary of materials and methods. For a detailed account, please refer to Supplementary Information.

### Human Brain Collection and Donor Characterization

Postmortem BA9 tissue was obtained from the NIH NeuroBioBank Brain Tissue Resource Center at Harvard University. TD cases were collected through the Tourette Association of America. The final cohort consisted of ten male individuals (five TD, five neurotypical controls), matched for age (controls: 25.0 ± 3.5 years; TD: 24.8 ± 5.9 years; P = 0.89) and PMI (controls: 20.6 ± 2.4 h; TD: 19.0 ± 4.4 h; P = 0.76). Diagnoses were confirmed by experienced clinicians. Controls were screened to exclude psychiatric disorders.

### Single-Nucleus RNA Sequencing

Nuclei were isolated by mechanical dissociation in neuroprotective buffer, purified through sucrose cushion centrifugation, and submitted for 10x Genomics library preparation and sequencing [30]. Raw data were processed using Cell Ranger with alignment to GRCh38. Quality control and downstream analyses were conducted in Seurat (version 4.1.0), with nuclei filtered for minimum 1,000 genes and maximum 5% mitochondrial content [31–34].

### Cell-Type Annotation and Differential Expression

Automated cell-type classification was performed using Azimuth against a custom BA9 reference generated from PsychENCODE single-nucleus data [35], validated by manual marker inspection. DEGs were identified using FindAllMarkers (FDR < 0.05, minimum 1.5-fold change). GO enrichment was performed using enrichR. Pseudotime analysis was conducted using Monocle3 [36, 37], with trajectories compared between groups using Wilcoxon rank-sum tests.

### Stress-Responsive and Cross-Disorder Analyses

DEGs were compared with dexamethasone-responsive gene sets [23] and a curated IEG reference [24]. Cross-disorder overlap with MDD and PTSD DLPFC signatures [23] was quantified using GeneOverlap [38].

### Integration with TD Genetic Architecture

BA9 DEGs were compared with gene sets from the TD GWAS meta-analysis [11], including eQTL/sQTL-linked genes, MAGMA gene-level associations (c-MAGMA, h-MAGMA, e-MAGMA), and brainSMR cell-type-specific associations.

### Cross-Regional Comparisons

BA9 DEGs were compared with published striatal snRNA-seq data from TD [7] using GeneOverlap [38] across all pairwise cell-type comparisons.

## ACKNOWLEDGMENTS

This study was partially supported by NINDS R21-NS125654. Postmortem human samples were obtained through the NIH NeuroBioBank from the Harvard Brain Tissue Resource Center/NIH NeuroBioBank. We acknowledge the Tourette Association of America (TAA) for spearheading, organizing and supporting the postmortem brain collection for patients with Tourette disorder. The authors also acknowledge the Cancer Genomics Shared Resource and HCI’s National Cancer Institute Center Support Grant (P30CA042014) and thank Dr. Brian Dalley and the staff for their assistance with the snRNA-seq experiments.

## DISCLOSURES

The authors declare no conflict of interest.

## DATA AND MATERIAL AVAILABILITY

All data needed to evaluate the conclusions in the paper are present in the paper and/or the Supplementary Materials. The data generated for snRNA-seq are available on Gene Expression Omnibus, GSE311334.

## Notes

### Competing Interest Statement

The authors have declared no competing interest.

### Summary of Updates

This version incorporates two key comparative analyses: (i) GWAS data from Tourette disorder (Strom et al., 2025) to examine genetic convergence with TD transcriptomic signatures, and (ii) snRNA-seq data from striatal tissue of TD patients (Wang et al., 2025) to assess cross-regional convergence between the DLPFC and striatum.

## REFERENCES

1. American Psychiatric Association, editor. Diagnostic and statistical manual of mental disorders: DSM-5-TR. Fifth edition, text revision. Washington, DC: American Psychiatric Association Publishing; 2022.

2. Leckman JF, King RA, Bloch MH. Clinical Features of Tourette Syndrome and Tic Disorders. J Obsessive Compuls Relat Disord. 2014;3:372–379.

3. Morand-Beaulieu S, Smith SD, Ibrahim K, Wu J, Leckman JF, Crowley MJ, et al. Electrophysiological signatures of inhibitory control in children with Tourette syndrome and attention-deficit/hyperactivity disorder. Cortex. 2022;147:157–168.

4. Felling RJ, Singer HS. Neurobiology of tourette syndrome: current status and need for further investigation. J Neurosci. 2011;31:12387–12395.

5. Kalanithi PSA, Zheng W, Kataoka Y, DiFiglia M, Grantz H, Saper CB, et al. Altered parvalbumin-positive neuron distribution in basal ganglia of individuals with Tourette syndrome. Proc Natl Acad Sci U S A. 2005;102:13307–13312.

6. Kataoka Y, Kalanithi PSA, Grantz H, Schwartz ML, Saper C, Leckman JF, et al. Decreased number of parvalbumin and cholinergic interneurons in the striatum of individuals with Tourette syndrome. J Comp Neurol. 2010;518:277–291.

7. Wang Y, Fasching L, Wu F, Suvakov M, Huttner A, Berretta S, et al. Interneuron Loss and Microglia Activation by Transcriptome Analyses in the Basal Ganglia of Tourette Disorder. Biological Psychiatry. 2025;98:260–270.

8. Lennington JB, Coppola G, Kataoka-Sasaki Y, Fernandez TV, Palejev D, Li Y, et al. Transcriptome Analysis of the Human Striatum in Tourette Syndrome. Biol Psychiatry. 2016;79:372–382.

9. Albin RL, Mink JW. Recent advances in Tourette syndrome research. Trends Neurosci. 2006;29:175–182.

10. Robertson MM, Eapen V, Singer HS, Martino D, Scharf JM, Paschou P, et al. Gilles de la Tourette syndrome. Nat Rev Dis Primers. 2017;3:16097.

11. Strom NI, Halvorsen MW, Grove J, Ásbjörnsdóttir B, Luðvígsson P, Thorarensen Ó, et al. Genome-Wide Association Study Meta-Analysis of 9619 Cases With Tic Disorders. Biol Psychiatry. 2025;97:743–752.

12. Barbey AK, Koenigs M, Grafman J. Dorsolateral prefrontal contributions to human working memory. Cortex. 2013;49:1195–1205.

13. Miller EK, Cohen JD. An integrative theory of prefrontal cortex function. Annu Rev Neurosci. 2001;24:167–202.

14. Peterson BS, Skudlarski P, Anderson AW, Zhang H, Gatenby JC, Lacadie CM, et al. A functional magnetic resonance imaging study of tic suppression in Tourette syndrome. Arch Gen Psychiatry. 1998;55:326–333.

15. Rae CL, Parkinson J, Betka S, Gouldvan Praag CD, Bouyagoub S, Polyanska L, et al. Amplified engagement of prefrontal cortex during control of voluntary action in Tourette syndrome. Brain Commun. 2020;2:fcaa199.

16. Joyce MKP, Uchendu S, Arnsten AFT. Stress and Inflammation Target Dorsolateral Prefrontal Cortex Function: Neural Mechanisms Underlying Weakened Cognitive Control. Biol Psychiatry. 2025;97:359–371.

17. Glausier JR, Datta D, Fish KN, Chung DW, Melchitzky DS, Lewis DA. Laminar Differences in the Targeting of Dendritic Spines by Cortical Pyramidal Neurons and Interneurons in Human Dorsolateral Prefrontal Cortex. Neuroscience. 2021;452:181–191.

18. Godar SC, Bortolato M. What makes you tic? Translational approaches to study the role of stress and contextual triggers in Tourette syndrome. Neurosci Biobehav Rev. 2017;76:123– 133.

19. Hoekstra PJ, Anderson GM, Limburg PC, Korf J, Kallenberg CGM, Minderaa RB. Neurobiology and neuroimmunology of Tourette’s syndrome: an update. Cell Mol Life Sci. 2004;61:886–898.

20. Conelea CA, Woods DW, Brandt BC. The impact of a stress induction task on tic frequencies in youth with Tourette Syndrome. Behav Res Ther. 2011;49:492–497.

21. Haber SN, Liu H, Seidlitz J, Bullmore E. Prefrontal connectomics: from anatomy to human imaging. Neuropsychopharmacol. 2022;47:20–40.

22. Arnsten AFT. Stress signalling pathways that impair prefrontal cortex structure and function. Nat Rev Neurosci. 2009;10:410–422.

23. Chatzinakos C, Pernia CD, Morrison FG, Iatrou A, McCullough KM, Schuler H, et al. Single-Nucleus Transcriptome Profiling of Dorsolateral Prefrontal Cortex: Mechanistic Roles for Neuronal Gene Expression, Including the 17q21.31 Locus, in PTSD Stress Response. Am J Psychiatry. 2023;180:739–754.

24. Wu YE, Pan L, Zuo Y, Li X, Hong W. Detecting Activated Cell Populations Using Single-Cell RNA-Seq. Neuron. 2017;96:313–329.e6.

25. Rădulescu A, Herron J, Kennedy C, Scimemi A. Global and local excitation and inhibition shape the dynamics of the cortico-striatal-thalamo-cortical pathway. Sci Rep. 2017;7:7608.

26. Gallopin T, Geoffroy H, Rossier J, Lambolez B. Cortical Sources of CRF, NKB, and CCK and Their Effects on Pyramidal Cells in the Neocortex. Cereb Cortex. 2006;16:1440–1452.

27. Chen P, Lou S, Huang Z-H, Wang Z, Shan Q-H, Wang Y, et al. Prefrontal Cortex Corticotropin-Releasing Factor Neurons Control Behavioral Style Selection under Challenging Situations. Neuron. 2020;106:301–315.e7.

28. Hu E, Demmou L, Cauli B, Gallopin T, Geoffroy H, Harris-Warrick RM, et al. VIP, CRF, and PACAP act at distinct receptors to elicit different cAMP/PKA dynamics in the neocortex. Cereb Cortex. 2011;21:708–718.

29. Georgiou C, Kehayas V, Lee KS, Brandalise F, Sahlender DA, Blanc J, et al. A subpopulation of cortical VIP-expressing interneurons with highly dynamic spines. Commun Biol. 2022;5:352.

30. Zheng GXY, Terry JM, Belgrader P, Ryvkin P, Bent ZW, Wilson R, et al. Massively parallel digital transcriptional profiling of single cells. Nat Commun. 2017;8:14049.

31. Butler A, Hoffman P, Smibert P, Papalexi E, Satija R. Integrating single-cell transcriptomic data across different conditions, technologies, and species. Nat Biotechnol. 2018;36:411– 420.

32. Romanov RA, Tretiakov EO, Kastriti ME, Zupancic M, Häring M, Korchynska S, et al. Molecular design of hypothalamus development. Nature. 2020;582:246–252.

33. Satija R, Farrell JA, Gennert D, Schier AF, Regev A. Spatial reconstruction of single-cell gene expression data. Nat Biotechnol. 2015;33:495–502.

34. Stuart T, Butler A, Hoffman P, Hafemeister C, Papalexi E, Mauck WM, et al. Comprehensive integration of single cell data. 2018.

35. Zintel T. Cellular Diversity in Human Subgenual Anterior Cingulate and Dorsolateral Prefrontal Cortex by Single-Nucleus RNA-sequencing. 2022.

36. Cao J, Spielmann M, Qiu X, Huang X, Ibrahim DM, Hill AJ, et al. The single-cell transcriptional landscape of mammalian organogenesis. Nature. 2019;566:496–502.

37. Trapnell C, Cacchiarelli D, Grimsby J, Pokharel P, Li S, Morse M, et al. The dynamics and regulators of cell fate decisions are revealed by pseudotemporal ordering of single cells. Nat Biotechnol. 2014;32:381–386.

38. Li Shen MS <Shenli SC. GeneOverlap. 2017.

