## Supplementary Information for "Tourette disorder features pervasive neuronal and glial transcriptional remodeling in the dorsolateral prefrontal cortex"

### SUPPLEMENTARY FIGURES

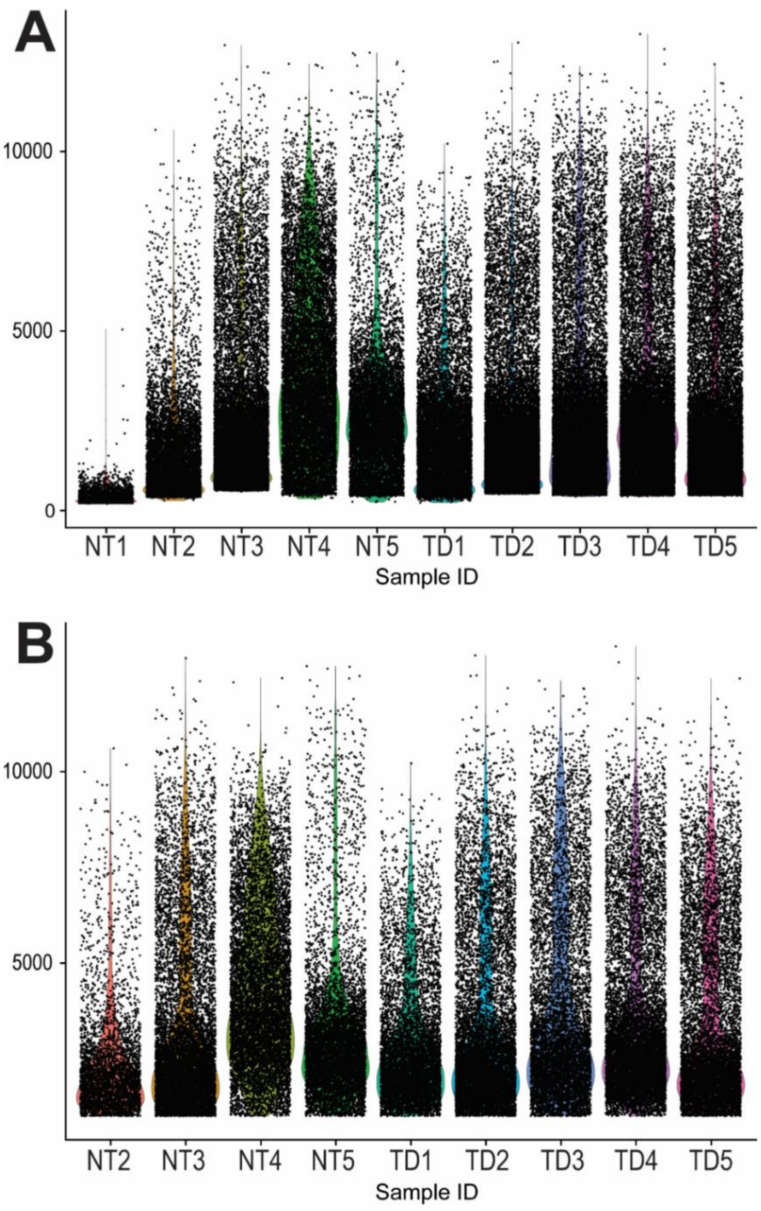


**Supplementary Figure 1:** Violin plots showing the number of detected features (nFeatures) per sample following initial processing with the Cell Ranger pipeline. (B) Violin plots of nFeatures for the same samples after quality control filtering, applying an nFeatures threshold >1,000 and a mitochondrial content cutoff <5%. Sample NT1 did not contain any nuclei meeting these criteria and was therefore excluded from downstream analyses; subsequent analyses were performed on samples NT2-NT5 and TD1-TD5.Abbreviations: NT, neurotypical; TD, Tourette Disorder.

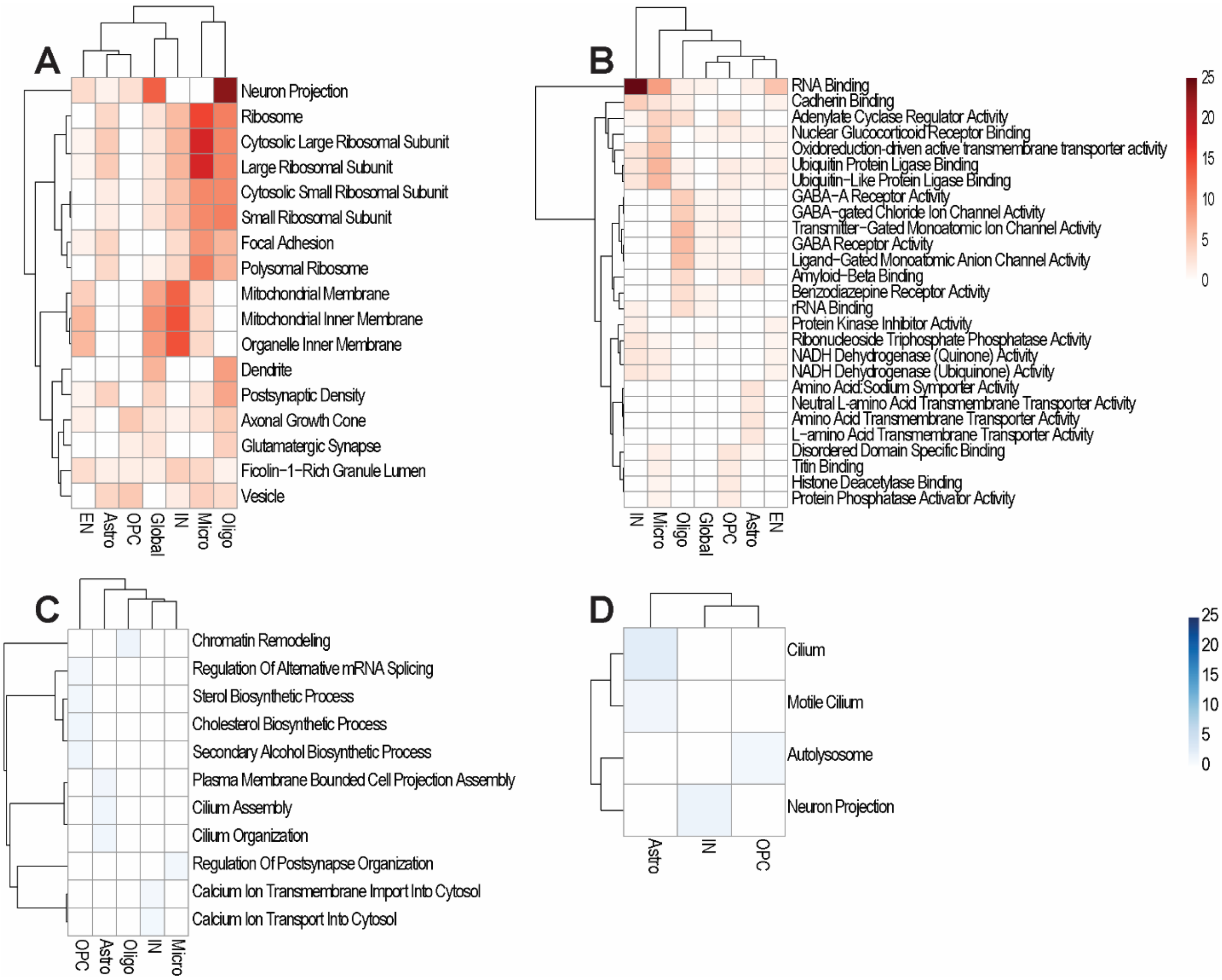


**Supplementary Figure 2.** Gene Ontology (GO) enrichment analyses for the comparison between neurotypical and TD samples. Related to Figure 1. (A) Upregulated cellular component terms; (B) upregulated molecular function terms; (C) downregulated biological process terms; and (D) downregulated cellular component terms. No significant GO terms were identified for downregulated molecular functions. Abbreviations: EN, excitatory neurons; IN, interneurons; Astro, astrocytes; Oligo, oligodendrocytes; OPC, oligodendrocyte precursor cell; Micro, microglia.

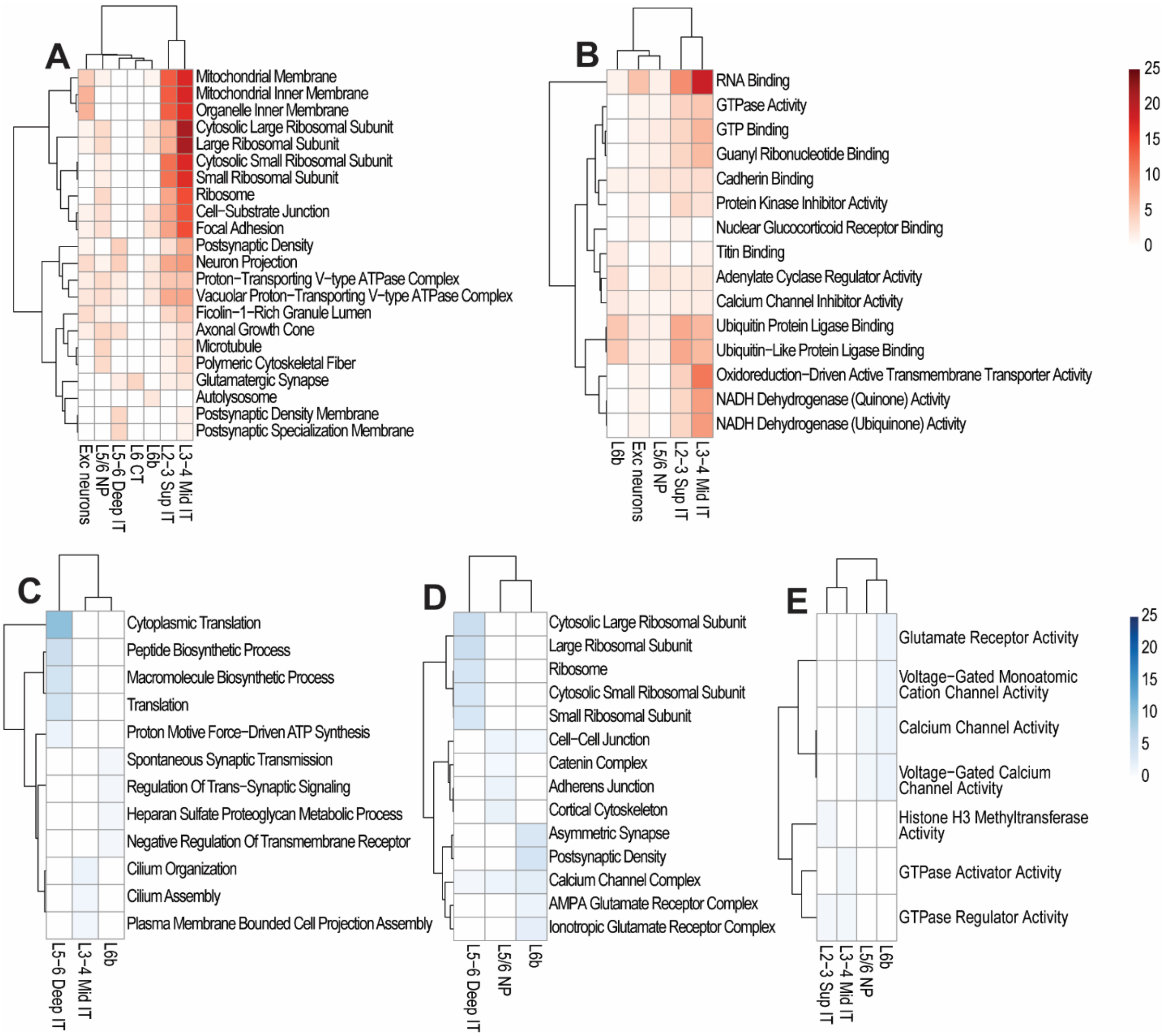


**Supplementary Figure 3.** Gene Ontology (GO) enrichment analyses for the comparison between neurotypical and TD samples for the excitatory neuron populations. Related to Figure 2. (A) Upregulated cellular component terms; (B) upregulated molecular function terms; (C) downregulated biological process terms; and (D) downregulated cellular component terms; (E) downregulated cellular molecular functions. Abbreviations: EN, excitatory neurons; L2-3 Sup IT, Layer 2-3 Superficial Intratelencephalic; L3-4 Mid IT, Layer 3-4 Middle Intratelencephalic; L5-6 Deep IT, Layer 5-6 Deep Intratelencephalic; L5/6 NP, Layer 5/6 Near-Projecting neurons; L6 CT, Layer 6 Corticothalamic; and L6b, Layer 6b.

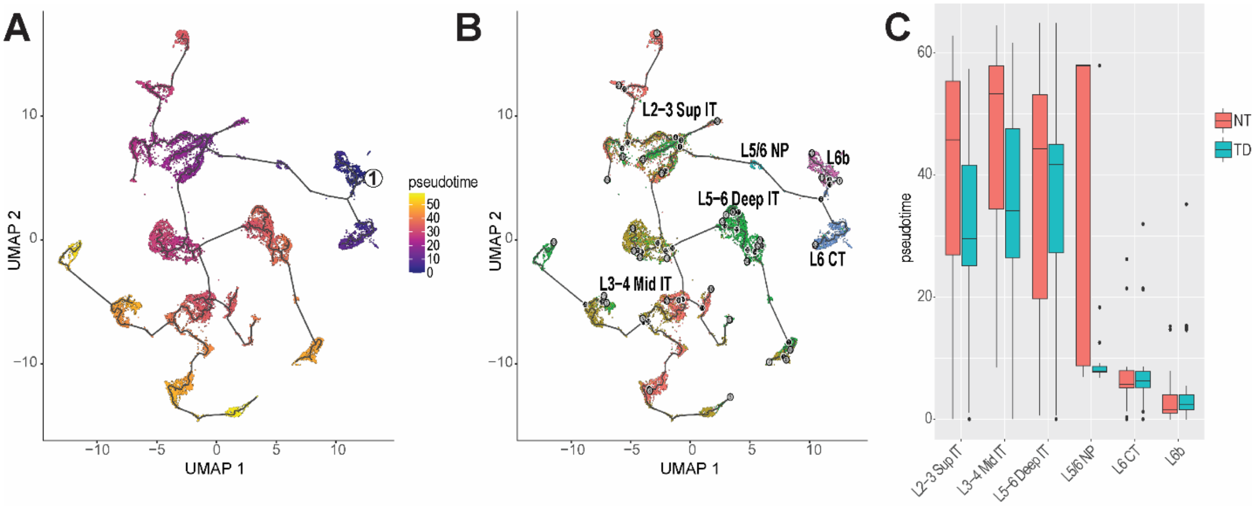


**Supplementary Figure 4.** Pseudotime trajectory analysis of excitatory neurons in neurotypical (NT, salmon) and Tourette disorder (TD, teal) samples. (A–B) UMAP embedding of excitatory neurons colored by pseudotime as inferred by Monocle3 (A) and annotated by excitatory neuronal subtypes (B). (C) Bar plots showing pseudotime distributions across excitatory neuronal subtypes. Abbreviations: L2-3 Sup IT, Layer 2-3 Superficial Intratelencephalic; L3-4 Mid IT, Layer 3-4 Middle Intratelencephalic; L5-6 Deep IT, Layer 5-6 Deep Intratelencephalic; L5/6 NP, Layer 5/6 Near-Projecting neurons; L6 CT, Layer 6 Corticothalamic; and L6b, Layer 6b.

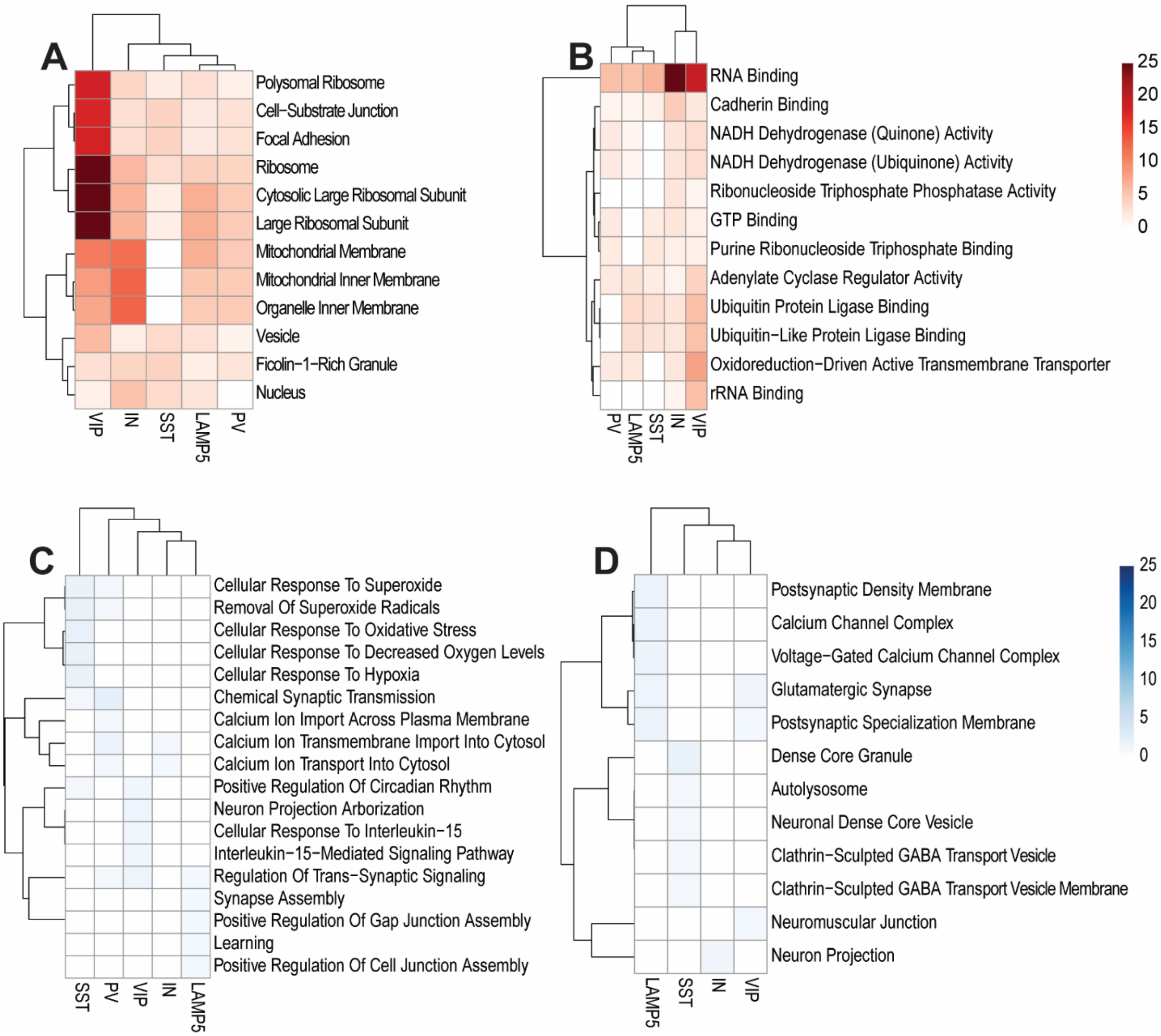


**Supplementary Figure 5.** Gene Ontology (GO) enrichment analyses for the comparison between neurotypical and TD samples for the interneuron populations. Related to Figure 3. (A) Upregulated cellular component terms; (B) upregulated molecular function terms; (C) downregulated biological process terms; and (D) downregulated cellular component terms. No significant GO terms were identified for downregulated molecular functions. Abbreviations: SST, somatostatin-positive; PV, parvalbumin-positive.

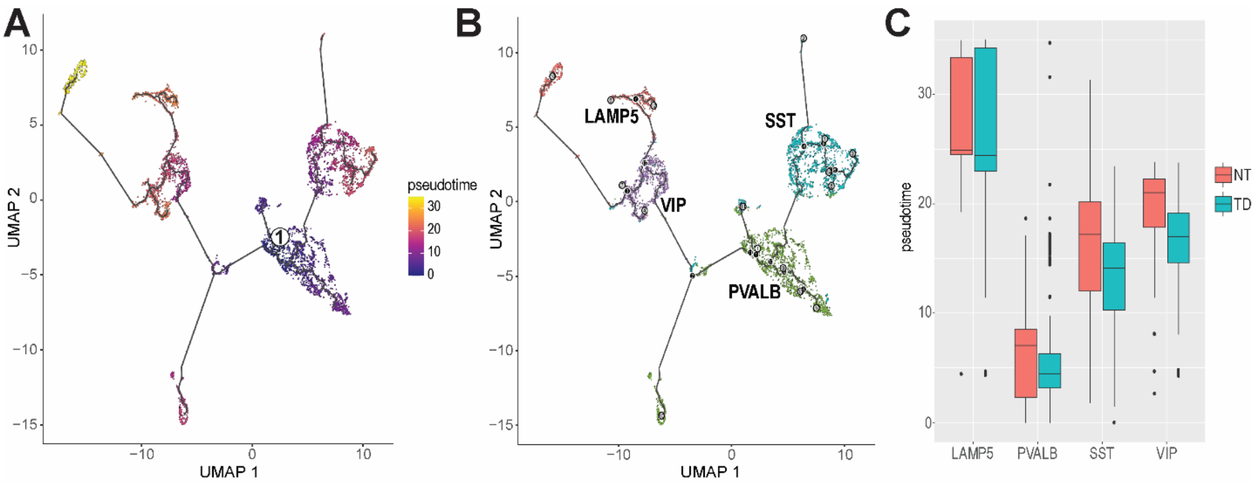


**Supplementary Figure 6.** Pseudotime trajectory analysis of interneurons in neurotypical (NT, salmon) and Tourette disorder (TD, teal) samples. (A–B) UMAP embedding of inhibitory neurons colored by pseudotime as inferred by Monocle3 (A) and annotated by interneuron subtypes (B). (C) Bar plots showing pseudotime distributions across inhibitory interneuron subclasses in NT and TD samples. Abbreviations: SST, somatostatin-positive; PV, parvalbumin-positive.

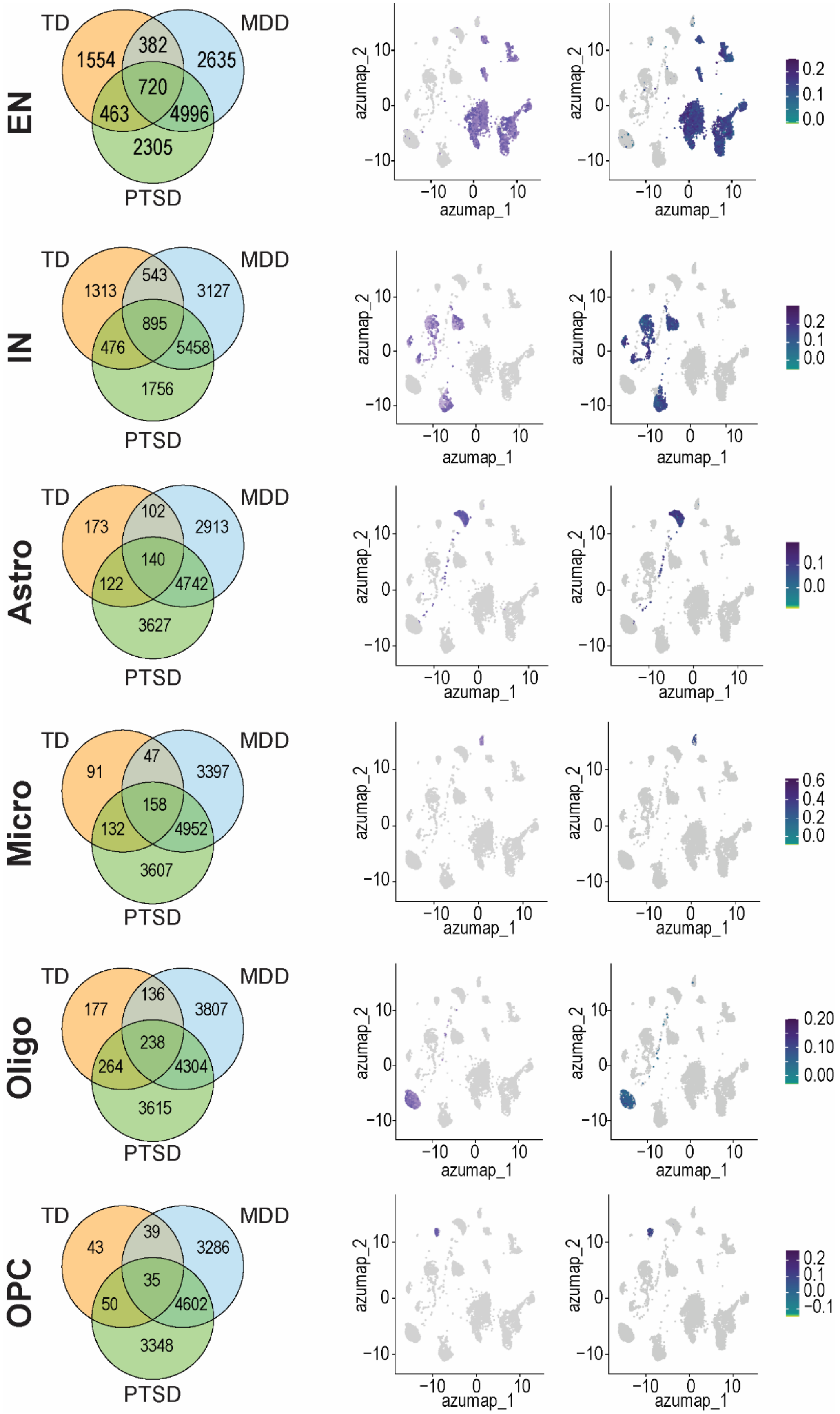
EN
TD
MDD
PTSD
1554
382
2635
720
463
4996
2305
IN
1313
543
3127
895
476
5458
1756
Astro
173
102
2913
140
122
4742
3627
Micro
91
47
3397
158
132
4952
3607
Oligo
177
136
3807
238
264
4304
3615
OPC
43
39
3286
35
50
4602
3348
azumap_1
azumap_2

**Supplementary Figure 7.** Cell-type–specific overlap between TD-, MDD-, and PTSD-associated transcriptional signatures and visualization of shared stress-responsive gene programs. Venn diagrams (left column) depict the intersection of upregulated genes identified in the DLPFC of individuals with TD and published single-nucleus transcriptomic datasets from major depressive disorder (MDD) and post-traumatic stress disorder (PTSD) [1]. Overlaps are shown separately for excitatory neurons (EN), interneurons (IN), astrocytes (astro), microglia (Micro), oligodendrocytes (Oligo), and oligodendrocyte precursor cells (OPC). Each diagram illustrates the number of DEGs unique to each condition as well as the genes shared across disorders, highlighting the substantial convergence of stress-related transcriptional programs between TD and these established stress-linked conditions.

The accompanying UMAP feature plots show the distribution and relative expression of the overlapping TD–MDD and TD–PTSD gene sets across the integrated DLPFC snRNA-seq atlas. Darker colors reflect higher module scores, indicating that stress-related transcriptional programs shared with MDD and PTSD are enriched in multiple neuronal and glial populations in TD. Overall, these results highlight a broad, multicellular convergence of stress- and glucocorticoid-responsive gene signatures throughout the DLPFC in TD.

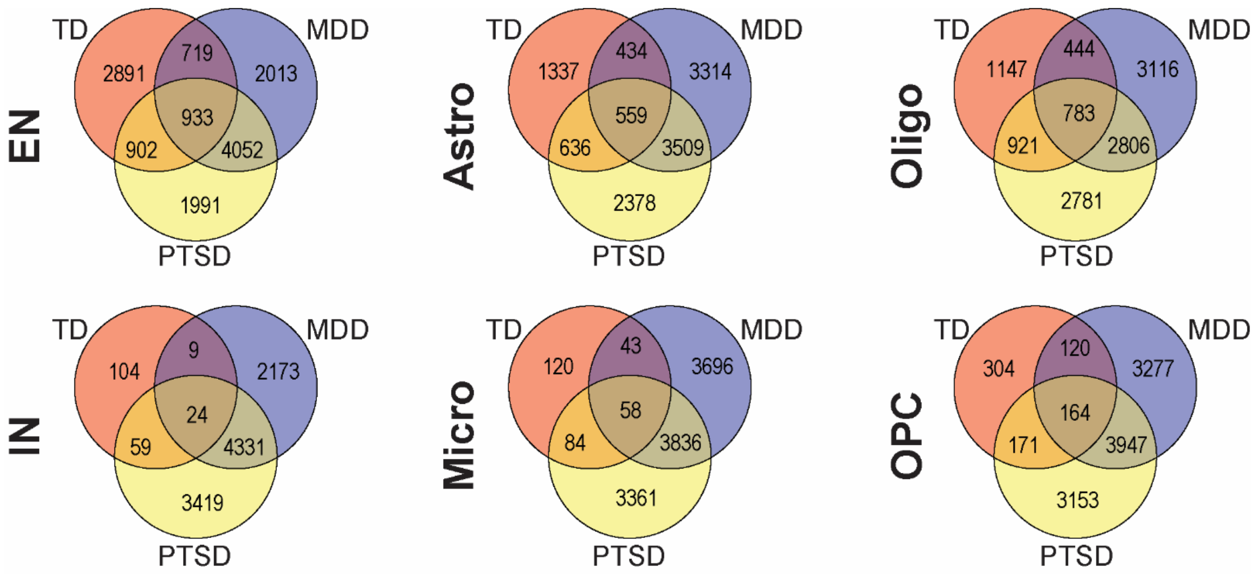


**Supplementary Figure 8.** Cell-type–specific Venn diagrams showing the intersection of downregulated genes identified in the DLPFC of individuals with Tourette disorder (TD) and those reported in published single-nucleus transcriptomic datasets from major depressive disorder (MDD) and post-traumatic stress disorder (PTSD) [1]. Each diagram indicates the number of DEGs unique to each condition, as well as the genes shared across disorders. Abbreviations: EN, excitatory neuron; IN, interneuron; Astro, astrocyte; Micro, microglia; Oligo, oligodendrocyte; OPC, oligodendrocyte precursor cell.

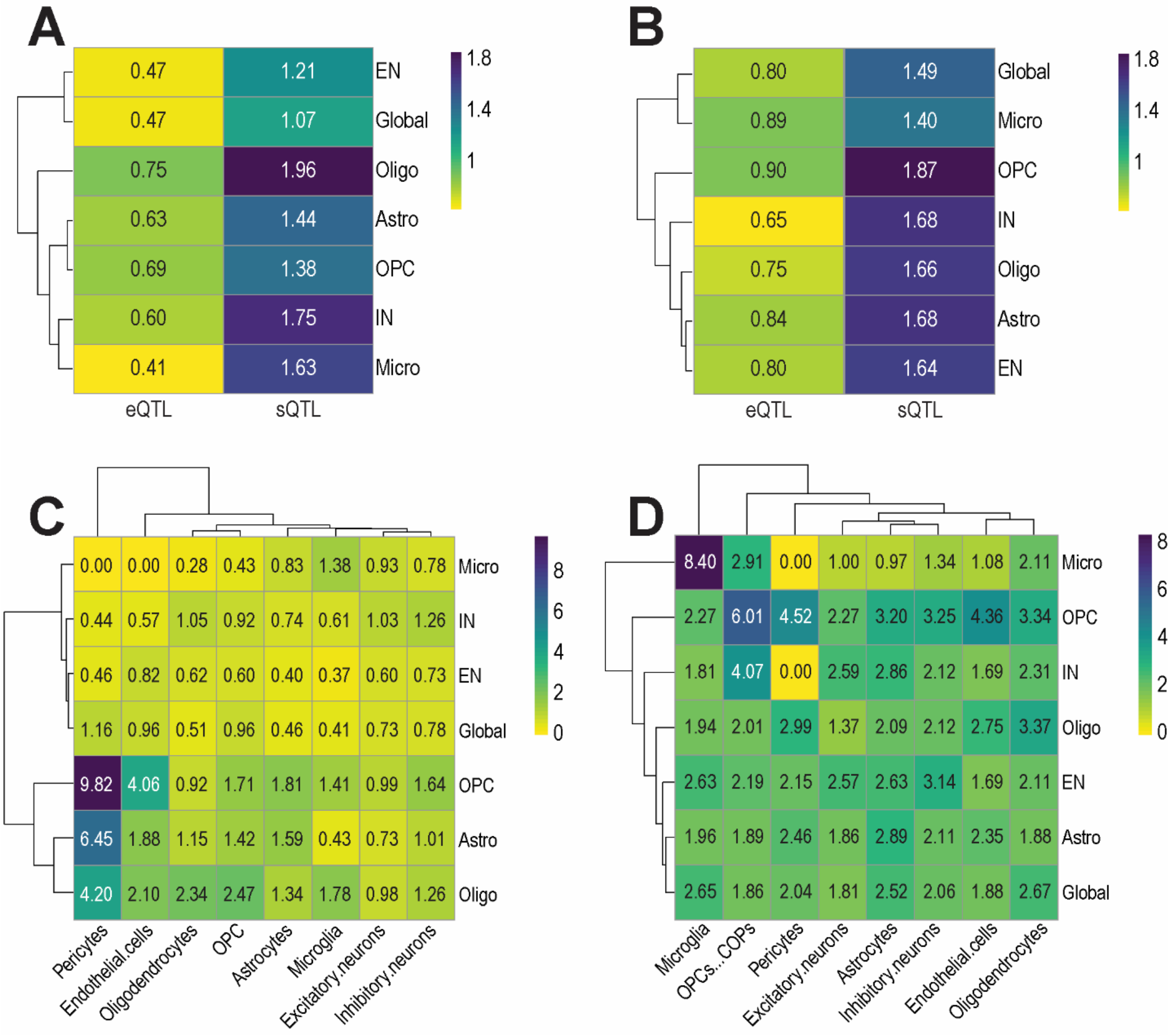


**Supplementary Figure 9.** Enrichment of eQTL-, sQTL-, and brainSMR-linked genes among differentially expressed genes. Heatmaps showing odds ratios (OR) for enrichment of eQTL- and sQTL-linked genes (list obtained by [2]) among upregulated (A) and downregulated DEGs (B) across cell types. Heatmaps showing OR for enrichment of brainSMR genes (list obtained by [2]) among upregulated (C) and downregulated DEGs (D) across cell types. Abbreviations: EN, excitatory neurons; IN, interneurons; Astro, astrocytes; Oligo, oligodendrocytes; OPC, oligodendrocyte precursor cell; Micro, microglia.

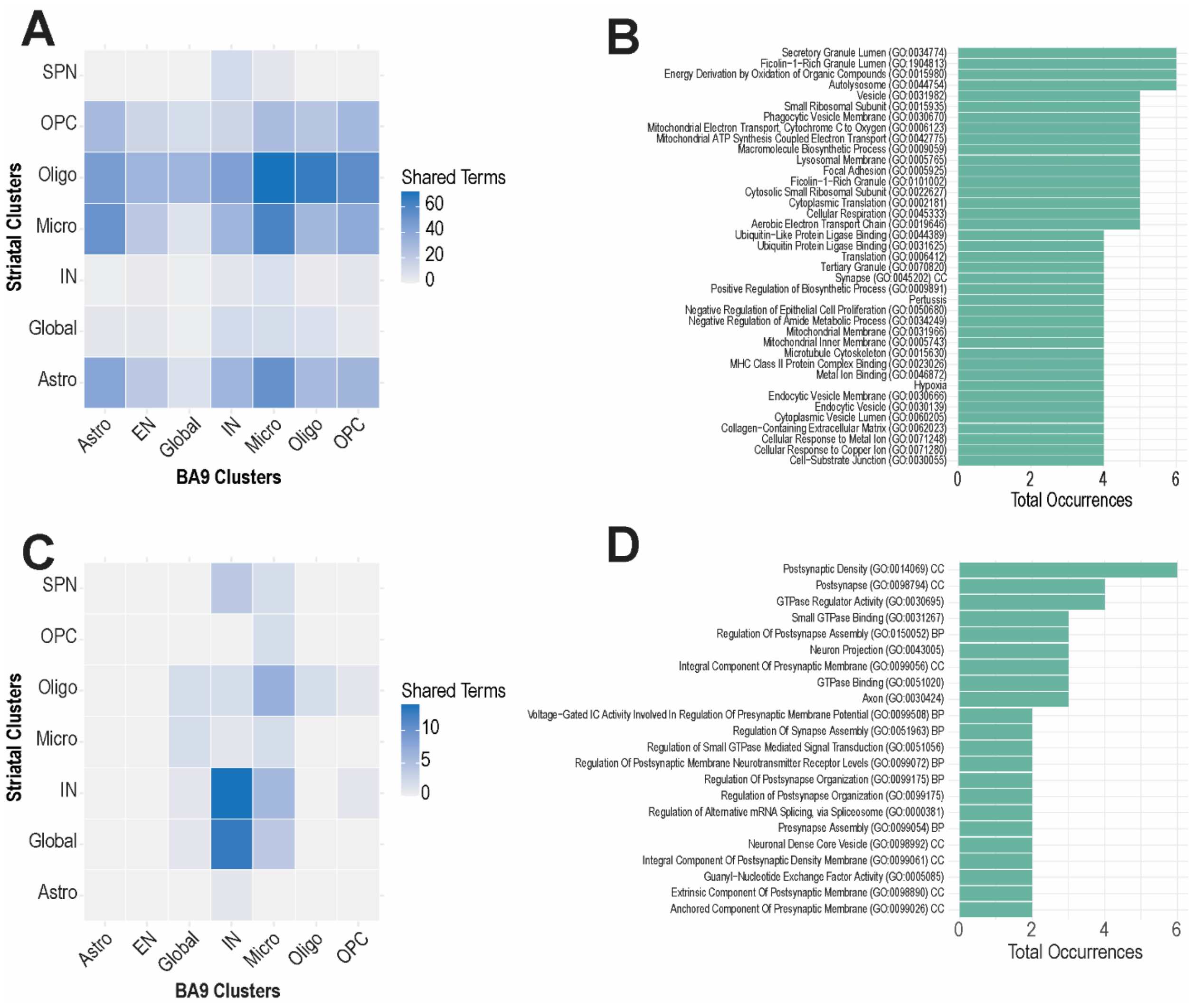


**Supplementary Figure 10.** Cluster-specific overlap between pathway enrichment results obtained from striatal samples in [3] and those generated in BA9, showing shared pathways among upregulated genes (A–B) and downregulated genes (C–D). Abbreviations: EN, Excitatory neuron; IN, interneuron; Astro, astrocyte; Micro, microglia; Oligo, oligodendrocyte; OPC, oligodendrocyte precursor cell; SPN, striatal projection neurons.

### SUPPLEMENTARY RESULTS

**Sample collection and cell-type identification.** Frozen BA9 samples were obtained from 10 male individuals, including 5 with TD and 5 age-matched neurotypical controls. Single-nucleus RNA sequencing (snRNA-seq) was performed on isolated nuclear fractions. Quality control filtering was applied using thresholds of >1,000 detected features and <5% mitochondrial content. One control sample (NT1) did not yield nuclei meeting these criteria and was excluded from downstream analyses (**Supplementary Fig. 1**). In total, 72,340 high-quality nuclei were retained for analysis (29,159 from controls; 43,181 from TD samples).

Unsupervised clustering identified six major cell populations (Figure 1A–B): excitatory neurons (23,276 nuclei; 30.8%), interneurons (7,015; 8.3%), astrocytes (9,657; 12.0%), oligodendrocytes (27,321; 36.4%), oligodendrocyte precursor cells (OPCs; 4,392; 6.1%), and microglia (3,093; 4.3%). In contrast to prior reports of interneuron depletion in the TD striatum [3–5], no significant differences in overall cell-type proportions were observed between TD and control samples (**Figure 1C**).

Differential expression analysis indicated widespread transcriptional changes across all major cell populations, with both the extent and direction of regulation differing by cell type. Interneurons exhibited the strongest directional bias, characterized predominantly by upregulated genes (3,227 up vs. 196 down), while microglia also showed a milder skew toward upregulation (733 DEGs: 428 up, 305 down). In contrast, excitatory neurons (8,564 DEGs: 3,119 up, 5,455 down), astrocytes (3,503 DEGs: 537 up, 2,966 down), oligodendrocytes (4,110 DEGs: 815 up, 3,295 down), and OPCs (926 DEGs: 167 up, 759 down) displayed a clear predominance of downregulated transcripts (**Supplementary Data 1-3**).

Despite variability in the overall direction of differential expression, GO enrichment analyses revealed a consistent functional signature across major cell types: significant enrichment was driven predominantly by upregulated genes, while downregulated gene sets showed minimal enrichment across categories (**Figure 1D**; **Supplementary Fig. 2**). The number of significant Gene Ontology (GO) terms (padj < 0.05) in the upregulated fraction, listed from most to least enriched for Biological Process, Cellular Component, and Molecular Function respectively, was: interneurons (126, 63, 12), OPCs (138, 10, 23), oligodendrocytes (106, 35, 26), microglia (89, 48, 17), astrocytes (77, 24, 16), and excitatory neurons (38, 20, 4), whereas downregulated terms were sparse or absent in most cell types.

GO enrichment analyses revealed a coherent cross-cell-type signature characterized by increased biosynthetic demand, enhanced proteostasis, and elevated mitochondrial and metabolic activity in TD brains, alongside cell type–specific adaptations in synaptic and signaling pathways. Overall, these findings point to a coordinated shift toward heightened translational and metabolic states across both neuronal and glial populations. At the global level, TD brains showed strong enrichment for *Translation* (GO:0006412), *Macromolecule Biosynthetic Process* (GO:0009059), and *Cytoplasmic Translation* (GO:0002181), accompanied by *Mitochondrial Translation* (GO:0032543) and *Mitochondrial Gene Expression* (GO:0140053). In excitatory neurons, upregulated genes were enriched for pathways related to protein synthesis and folding, including *Translation* (GO:0006412), *Macromolecule Biosynthetic Process* (GO:0009059), and *'De Novo' Post-Translational Protein Folding* (GO:0051084). Prominent enrichment was also observed for mitochondrial organization and protein targeting processes. In interneurons, upregulated genes were strongly enriched for pathways related to protein synthesis and gene expression, including *Translation* (GO:0006412), *Cytoplasmic Translation* (GO:0002181), *Macromolecule Biosynthetic Process* (GO:0009059), *and Gene Expression* (GO:0010467). Additional enrichment was observed in intracellular trafficking processes, such *as Intracellular Protein Transport* (GO:0006886) and *Endoplasmic Reticulum to Golgi Vesicle-Mediated Transport* (GO:0006888). Notably, interneurons also showed prominent enrichment for mitochondrial and metabolic pathways, including *Oxidative Phosphorylation* (GO:0006119), *Mitochondrial Translation* (GO:0032543), and *Mitochondrial Gene Expression* (GO:0140053). In astrocytes, upregulated genes were significantly enriched for processes related to proteostasis and cellular stress responses, including *Response to Unfolded Protein* (GO:0006986) and *Chaperone Cofactor-Dependent Protein Refolding* (GO:0051085), alongside increased biosynthetic and translational activity (*Macromolecule Biosynthetic Process*, GO:0009059; *Cytoplasmic Translation*, GO:0002181). Enrichment was also observed for vascular and transport-related functions, as well as metal ion response pathways. In microglia, upregulated genes were enriched for pathways related to protein synthesis and overall metabolic activity, including *Cytoplasmic Translation* (GO:0002181), *Translation* (GO:0006412), *Macromolecule Biosynthetic Process* (GO:0009059), *Gene Expression* (GO:0010467), and *Protein Metabolic Process* (GO:0019538). Strong enrichment was also observed for mitochondrial and energy metabolism pathways, such as *Cellular Respiration* (GO:0045333), *Energy Derivation by Oxidation of Organic Compounds* (GO:0015980), *Aerobic Electron Transport Chain* (GO:0019646), and *Mitochondrial ATP Synthesis Coupled Electron Transport* (GO:0042775). In oligodendrocytes, upregulated genes were enriched for pathways related to protein synthesis and gene expression, including *Cytoplasmic Translation* (GO:0002181), *Translation* (GO:0006412), *Macromolecule Biosynthetic Process* (GO:0009059), and *Gene Expression* (GO:0010467). In addition, these cells showed notable enrichment for synaptic and neurotransmission-related processes, such as *Chemical Synaptic Transmission* (GO:0007268), *Synaptic Transmission, GABAergic* (GO:0051932), *Gamma-Aminobutyric Acid Signaling Pathway* (GO:0007214), and *Synaptic Vesicle Exocytosis* (GO:0016079).

In OPCs, upregulated genes were enriched for pathways related to synaptic modulation and neuronal connectivity, including *Positive Regulation of Synaptic Transmission*

(GO:0050806) and *Axon Guidance* (GO:0007411). These cells also showed enrichment for proteostasis-related processes, such as *Response to Unfolded Protein* (GO:0006986), *Chaperone-Mediated Protein Complex Assembly* (GO:0051131), and *Chaperone Cofactor-Dependent Protein Refolding* (GO:0051085) (**Figure 1D** and **Supplementary Data 1**).

**Layer-specific transcriptional remodeling of excitatory neurons.**

To assess whether transcriptional alterations in TD were uniformly distributed across cortical layers or instead reflected layer-specific effects, we first stratified excitatory neurons into six transcriptionally distinct, layer-resolved subtypes (**Figure 2A–B**): L2–3 Superficial Intratelencephalic (IT), L3–4 Middle IT, and four deep-layer populations (L5–6 Deep IT, L5/6 Near-Projecting [NP], L6 Corticothalamic [CT], and L6b neurons). The relative proportions of these subtypes were comparable between TD and control samples (**Figure 2C**), indicating that disease-associated changes are not driven by shifts in layer-specific cell abundance.

Despite preserved composition, pronounced layer-specific transcriptional remodeling was observed (**Supplementary Data S4–6**). Deep-layer populations showed a predominance of upregulated genes, including L5/6 Deep IT (3,433 DEGs: 2,291 up, 1,142 down), L5/6 NP (373: 284 up, 89 down), and L6 CT (156: 93 up, 63 down). In contrast, superficial and middle layers were characterized by a strong bias toward downregulation, as observed in L2–3 Superficial IT (8,150: 1,703 up, 5,447 down), L3–4 Middle IT (5,986: 1,669 up, 4,317 down), and L6b neurons (661: 210 up, 451 down). This bidirectional pattern aligns with the functional architecture of cortical circuits, whereby superficial IT neurons primarily mediate corticocortical communication, middle-layer neurons constitute the principal thalamocortical input zone, and deep-layer neurons project to subcortical targets such as the striatum and thalamus [6].

Analysis of excitatory neuron subtypes revealed a coordinated, layer-dependent reprogramming in TD, with enhanced metabolic and translational activity in superficial neurons and a more complex remodeling in deep layers, combining synaptic upregulation with reduced biosynthetic and metabolic activity (**Figure 2D**, **Supplementary Fig. 3** and **Supplementary Data S4–6**). In superficial and middle excitatory neurons (L2–3 and L3–4 IT populations), upregulated genes converged on a shared signature of enhanced protein synthesis and metabolic activity, including *Translation* (GO:0006412), *Cytoplasmic Translation* (GO:0002181), *Macromolecule Biosynthetic Process* (GO:0009059), and *Gene Expression* (GO:0010467). This was accompanied by strong enrichment of mitochondrial and bioenergetic pathways, such as *Cellular Respiration* (GO:0045333), *Oxidative Phosphorylation* (GO:0006119), *Proton Motive Force-Driven ATP Synthesis* (GO:0015986), *Proton Motive Force-Driven Mitochondrial ATP Synthesis* (GO:0042776), and *Mitochondrial ATP Synthesis Coupled Electron Transport* (GO:0042775), with additional contributions from *Aerobic Respiration* (GO:0009060) and

*Proton Transmembrane Transport* (GO:1902600). Together, these findings indicate a coordinated upregulation of translational and mitochondrial processes across superficial and middle cortical layers, with only minor variations reflecting differences in protein metabolism and respiratory chain activity. In L5/6 NP neurons, upregulated genes were enriched for processes related to protein synthesis and intracellular transport, including *Cytoplasmic Translation* (GO:0002181), *Protein Exit From Endoplasmic Reticulum* (GO:0032527), and *Endoplasmic Reticulum to Cytosol Transport* (GO:1903513). Prominent enrichment was also observed for pathways regulating ion homeostasis and organelle acidification, such as *Proton Transmembrane Transport* (GO:1902600), *Golgi Lumen Acidification* (GO:0061795), *Endosomal Lumen Acidification* (GO:0048388), *Vacuolar Acidification* (GO:0007035), and *Intracellular pH Reduction* (GO:0051452). In L5/6 deep IT neurons, upregulated genes were enriched for pathways related to synaptic organization and neuronal development, including *Synapse Organization* (GO:0050808), *Axonogenesis* (GO:0007409), and *Nervous System Development* (GO:0007399). These cells also showed enrichment for processes governing neuronal structure and connectivity, such as *Negative Regulation of Neuron Projection Development* (GO:0010977) and *Negative Regulation of Cell Projection Organization* (GO:0031345), alongside prominent involvement of synaptic signaling and plasticity pathways, including *Modulation of Chemical Synaptic Transmission* (GO:0050804), *Synaptic Transmission, Glutamatergic* (GO:0035249), and *Regulation of Neuronal Synaptic Plasticity* (GO:0048168). Conversely, downregulated genes in this population were enriched for pathways related to protein synthesis and cellular bioenergetics, including *Cytoplasmic Translation* (GO:0002181), *Translation* (GO:0006412), *Macromolecule Biosynthetic Process* (GO:0009059), and *Proton Motive Force-Driven ATP Synthesis* (GO:0015986), indicating a relative reduction in translational and metabolic activity. In L6 CT neurons, upregulated genes were enriched for pathways related to synapse assembly and presynaptic organization, including *Regulation of Synapse Assembly* (GO:0051963), *Presynapse Assembly* (GO:0099054), *Presynaptic Membrane Assembly* (GO:0097105), and *Presynaptic Membrane Organization* (GO:0097090). Additional enrichment was observed in regulatory processes, such as *Regulation of Presynapse Assembly* (GO:1905606) and *Regulation of Presynapse Organization* (GO:0099174), as well as broader synaptic architecture pathways including *Synapse Organization* (GO:0050808). In L6b neurons, upregulated genes were enriched for pathways related to calcium signaling and ion homeostasis, as well as broader processes such as *Regulation of Calcium Ion Transmembrane Transporter Activity* (GO:1901019). These cells also showed enrichment for signaling and plasticity-related functions, including *Regulation of Cell Communication by Electrical Coupling* (GO:0010649) and *Long-Term Synaptic Potentiation* (GO:0060291).

To determine whether these transcriptional differences were associated with alterations in neuronal maturation, we performed pseudotime analysis within excitatory neurons. TD cells were predominantly enriched at earlier and intermediate pseudotime states, whereas neurons from neurotypical samples progressed toward more advanced maturation stages (**Supplementary Fig. 4**; **Supplementary Data S7**). Together, these findings suggest that TD is characterized by coordinated, layer-dependent transcriptional remodeling coupled with a relative shift toward less mature excitatory neuronal states, consistent with its classification as a neurodevelopmental disorder with an age-dependent trajectory.

**Widespread transcriptional activation in interneuron populations.** We next assessed differential gene expression across interneuron subtypes (**Figure 3**). Two major classes were identified: MGE-derived interneurons, defined by LHX6 expression and comprising PV+ (n = 2,140; 34.0%) and SST+ (n = 1,927; 30.8%) populations, and CGE-derived interneurons, marked by PROX1 and subdivided into VIP+ (n = 1,060; 18.4%) and LAMP5+ (n = 888; 16.8%) cells (**Figure 3A-B**). No significant differences in subtype proportions were observed between TD and control groups (**Figure 3C**).

Gene Ontology enrichment analyses (**Figure 3D**, **Supplementary Fig.5**, and **Supplementary Data S8-10**) further revealed a shared functional signature across PV+, VIP+, and LAMP5+ interneurons, characterized by increased protein synthesis and metabolic activity, including *Cytoplasmic Translation* (GO:0002181), *Translation* (GO:0006412), *Macromolecule Biosynthetic Process* (GO:0009059), and mitochondrial pathways such as *Oxidative Phosphorylation* (GO:0006119), *Cellular Respiration* (GO:0045333), *Aerobic Electron Transport Chain* (GO:0019646), and *Mitochondrial ATP Synthesis Coupled Electron Transport* (GO:0042775). This convergent profile suggests a common state of heightened biosynthetic and energetic demand across these interneuron populations, with subtype-specific nuances, PV+ neurons showing a stronger bias toward mitochondrial ATP production, and VIP+/LAMP5+ neurons exhibiting additional enrichment in ribosome biogenesis and gene expression pathways.

In contrast, SST+ interneurons displayed a distinct enrichment profile dominated by proteostasis and post-translational regulation, including *Regulation of Protein Ubiquitination* (GO:0031396), *Protein Stabilization* (GO:0050821), *Regulation of Proteasomal Ubiquitin-Dependent Protein Catabolic Process* (GO:0032434), and *'De Novo' Post-Translational Protein Folding* (GO:0051084), along with stress response pathways such as *Cellular Response to Heat* (GO:0034605).

Consistent with excitatory neuron findings, pseudotime analysis revealed that TD interneurons were preferentially enriched at earlier and intermediate maturation states, whereas control neurons progressed toward more advanced stages (**Supplementary Fig. 6; Supplementary Data S11**).

**Enrichment for glucocorticoid-responsive and immediate early gene modules.**

To evaluate whether the transcriptional alterations observed in the DLPFC of individuals with TD reflect broader stress-related molecular programs, we compared the DEGs identified in our dataset with published gene sets describing glucocorticoid-responsive modules [1]. In that study, dexamethasone exposure of iPSC-derived neurons yielded a well-defined set of corticosteroid-induced genes. Using these dexamethasone-responsive gene lists as a reference, we found a robust transcriptional enrichment across all major DLPFC cell populations in TD (**Figure 4A** and **Supplemental Data S12**), indicating marked upregulation of corticosteroid-linked molecular pathways.

Cell-type-specific analyses revealed distinct yet convergent patterns of glucocorticoid-responsive gene induction. In neuronal populations, both excitatory neurons and interneurons displayed increased expression of canonical immediate early genes (IEGs) together with metabolic and extracellular matrix regulators consistent with activity-dependent modulation and potential perisynaptic restructuring. Among non-neuronal populations, oligodendrocytes exhibited the strongest enrichment amongst the non-neuronal populations, with upregulation of osmotic and stress-adaptation genes such as *SLC5A3* and *HSPB1*, pointing to alterations in white-matter homeostasis. Astrocytes showed elevations in growth factor–associated transcripts (*IGFBP7*), immediate early genes (*FOS*, *FOSB*), and cell-cycle regulators (*CKS2*), consistent with a reactive transcriptional phenotype. Microglia showed significant upregulation of stress-response machinery such as *HSP90AB1* and multiple DNA replication factors (*TOP2A*, *MELK*, *RRM2*), suggesting enhanced activation or remodeling states (**Supplemental Data S12**). We then examined whether upstream hypothalamic-pituitary-adrenal (HPA) axis signaling may contribute to these glucocorticoid-associated signatures by interrogating expression of *CRH*, the gene encoding corticotropin-releasing factor. Expression of this gene was significantly upregulated across interneurons ($log_2FC = 0.96$, padj = $3.3 \times 10^{-13}$), with the strongest induction in PV+ ($log_2FC = 1.35$, padj = $4.5 \times 10^{-4}$) and VIP+ cells (log2FC = 1.11, padj = $1.8 \times 10^{-7}$), where over half of TD nuclei expressed detectable transcript versus ~30% in controls.

Leveraging the same study, we also analyzed the convergent molecular abnormalities between TD and MDD and PTSD [1]. Upon identification of the reference datasets, we first confirmed that cell-type cluster assignments were comparable across studies (**Supplementary Table 1**).

We observed substantial overlap between TD-associated DEGs and the shared MDD-PTSD gene set across multiple cortical cell populations (**Supplementary Figure 7-8** and **Supplemental Data S13**). The largest intersections of upregulated genes were found in excitatory neurons and interneurons, where considerable proportions of TD-overexpressed genes also appeared in both stress-related disorders. Although the absolute number of shared genes was smaller in non-neuronal cell types, consistent enrichment was nevertheless evident in astrocytes, microglia, oligodendrocytes, and OPCs. Notably, UMAP projections confirmed that this shared MDD–PTSD signature was selectively enriched within the same TD-upregulated clusters, reinforcing the spatial and transcriptional specificity of this convergence (**Supplementary Fig. 7**). These findings suggest that the TD transcriptional landscape recapitulates key components of stress-associated molecular dysregulation across both neuronal and glial compartments.

Taken together, these analyses indicate that the DLPFC transcriptional architecture in TD is characterized by broad upregulation of genes previously implicated in stress-related psychopathology. The convergence across disorders, cell types, and analytic modalities supports the interpretation that chronic or dysregulated stress-adaptation mechanisms may represent a significant component of TD pathophysiology, particularly within excitatory and inhibitory cortical neurons.

Beyond transcription factor–driven stress modules, we next assessed immediate early gene activity as a complementary, real-time indicator of cellular activation state. Using a curated IEG reference list [7], we identified strong enrichment across multiple TD cell populations, with the most pronounced activation signatures in microglia, followed by astrocytes and OPCs (**Supplementary Table 2** and **Supplementary Data S14**). A consistent pattern emerged across glial and neuronal compartments for the orphan nuclear receptor family genes *NR4A1*, *NR4A2*, and *NR4A3*, which regulate inflammatory responses, metabolic adaptation, and cellular stress resilience. Additional stress- and activity-associated transcripts, including *CCL2* in microglia and the IEGs *FOS*, *FOSB*, and *EGR1* across both neuronal and glial populations, were similarly elevated (**Figure 5B** and **Supplemental Data S14**). Because IEG expression can be influenced by perimortem conditions, we assessed potential confounds. PMI did not differ between groups and showed no significant correlation with IEG expression at the donor level (all padj > 0.45) or with cell-type-specific IEG module scores (all padj > 0.70). Single-nucleus PMI correlations were predominantly negative, opposite to what would be expected if PMI were inflating IEG signal. The IEG upregulation was group-specific and cell-type-patterned, whereas agonal artifacts would produce diffuse, nonspecific induction.

Together, these findings demonstrate that the DLPFC in TD exhibits coordinated activation of glucocorticoid-responsive transcriptional modules and immediate early gene networks. The convergence of corticosteroid-induced signatures with activity-dependent gene enrichment across neuronal and glial compartments supports the presence of a generalized hyperactivated cortical state in TD and reinforces the involvement of stress-adaptation pathways in its underlying pathophysiology.

**TD-associated transcriptional changes converge with the genetic architecture of TD.** The introduction noted that the largest TD GWAS meta-analysis identified polygenic risk enrichment in BA9-expressed genes [2]. Having characterized the transcriptional landscape of the TD DLPFC, including the stress-responsive programs described above, we next evaluated whether these alterations intersect with the common-variant genetic architecture of TD.

We compared BA9 DEGs with gene sets derived from the Strom et al. (2025) GWAS meta-analysis [2]. All three genes reaching significance in the GWAS gene-based analyses were differentially expressed in our dataset: *BCL11B* (log2FC = +0.23, padj = $5.4 \times 10^{-32}$) and *NDFIP2* (log2FC = +0.19, padj = $1.2 \times 10^{-14}$) were upregulated, while *RBM26* (log2FC = -0.16, padj = $6.9 \times 10^{-74}$) was downregulated.

Overlap with eQTL- and sQTL-linked genes revealed a striking dissociation (**Supplementary Figure 9A-B** and **Supplementary Data S15**): DEGs were uniformly depleted among eQTL-linked genes (upregulated: ORs 0.41-0.75; downregulated: ORs 0.65-0.90), yet consistently enriched among sQTL-linked genes (upregulated: ORs 1.07-1.96; downregulated: ORs 1.40-1.87), suggesting that TD risk variants preferentially influence splicing rather than expression levels of transcriptionally altered genes.

MAGMA analyses using conventional (c-MAGMA), chromatin interaction-informed (h-MAGMA), and eQTL-informed (e-MAGMA) approaches (**Supplementary Data S15**) revealed enrichment exclusively among upregulated DEGs under c-MAGMA and h-MAGMA, concentrated in OPCs, microglia, oligodendrocytes, and inhibitory neurons (ORs 1.27-1.97). e-MAGMA replicated this signal but additionally revealed broad enrichment among downregulated DEGs across most cell types (ORs 1.24-1.45), likely reflecting broadly expressed regulatory variants detectable only through eQTL-based mapping.

BrainSMR analyses revealed a further asymmetry (**Supplementary Figure 9C-D** and **Supplementary Data S16**). For upregulated DEGs, enrichment was concentrated in the oligodendrocyte lineage (with GWAS-implicated oligodendrocyte genes: OR = 2.34, P = $5.5 \times 10^{-8}$; OPC/COP genes: OR = 2.47, P = $4.6 \times 10^{-4}$). For downregulated DEGs, broad enrichment was observed across nearly all cell types (microglia: OR = 8.40; OPCs: OR = 6.01; oligodendrocytes: OR = 3.37; excitatory neurons: OR = 2.57; astrocytes: OR = 2.89; all $P < 10^{-11}$), except inhibitory neurons (OR = 2.12, P = 0.072), consistent with a genetically constrained transcriptional landscape.

**Cross-regional comparison with striatal transcriptomics reveals convergent glial programs and divergent neuronal pathology.** Finally, to determine whether the transcriptional changes in the DLPFC reflect region-specific or shared pathological processes across CSTC circuitry, we compared our BA9 DEGs with published striatal snRNA-seq data from TD [3]. Gene-level overlap analyses were performed across all pairwise cell-type comparisons, separately for upregulated and downregulated genes.

For upregulated genes, the strongest cross-regional convergence was observed among glial populations. Cell-type-matched comparisons revealed particularly high enrichment for microglia (OR = 20.2, P = $2.9 \times 10^{-12}$), OPCs (OR = 14.3, P = $8.5 \times 10^{-9}$), and astrocytes

(OR = 5.1, P = 9.4 x $10^{-17}$). Striatal oligodendrocytes emerged as a broader convergence hub, sharing upregulated genes with BA9 oligodendrocytes (74 genes, OR = 10.4, P = 5.5 x $10^{-43}$), BA9 microglia (61 genes, OR = 16.7, P = 8.3 x $10^{-47}$), and BA9 excitatory neurons (118 genes, OR = 4.7, P = 1.3 x $10^{-31}$). Comparison of independently derived GO enrichment results across the two cohorts revealed substantial functional convergence, with 76 overlapping GO terms in oligodendrocytes, 63 in microglia, 45 in astrocytes, and 34 in OPCs (**Supplementary Fig. 10A-B and Supplementary Data S17-18**).

For downregulated genes, striatal and BA9 interneurons exhibited significant overlap (OR = 5.7, P = 3.2 x $10^{-9}$), as well as microglia (OR = 18.2, P = 3.5 x $10^{-11}$), oligodendrocytes (OR = 3.2, P = 3.3 x 10-$^{105}$), and OPCs (OR = 39.4, P = 6.6 x $10^{-4}$). Cross-cohort comparison of GO enrichment results for downregulated genes revealed more limited functional convergence, with overlapping terms concentrated in inhibitory neurons (14 terms) and only 2 shared terms each in microglia and oligodendrocytes (**Supplementary Figure 10B-C** and **Supplementary Data S19-S20**).

TD shows a coordinated, cell type–specific reprogramming across brain regions, with glial populations adopting a broadly activated, metabolically and immunologically engaged state, while interneurons exhibit a distinct and opposing signature characterized by preserved vesicular machinery but widespread disruption of synaptic structure and inhibitory function. Cluster-resolved analyses integrating both up- and downregulated GO terms are presented below, organized by cell population, starting with glial populations and concluding with interneurons.

The most strongly enriched shared pathways in astrocytes fell into five broad functional categories. First, neurovascular and structural remodeling was highlighted by enrichment of *astrocyte end foot* (OR = 24.25), *focal adhesion* (OR = 2.93), *cell-substrate junction* (OR = 2.86), and *collagen-containing extracellular matrix* (OR = 2.04), consistent with reactive astrocytic remodeling of the neurovascular interface. Second, lipid and lipoprotein metabolism was represented by *high-density lipoprotein particle* (OR = 9.73), *low-density lipoprotein particle receptor binding* (OR = 9.62), *triglyceride-rich plasma lipoprotein particle* (OR = 9.11), and *very-low-density lipoprotein particle* (OR = 9.11), suggesting dysregulation of astrocytic lipid homeostasis across both regions. Third, oxidative and hypoxic stress responses were enriched, including *cellular response to hypoxia* (OR = 3.66), *cellular response to decreased oxygen levels* (OR = 4.19), *cellular response to metal ion* (OR = 4.85), and *cellular response to copper ion* (OR = 19.98), pointing to a shared metabolic stress signature. Fourth, vesicular and endocytic trafficking pathways were consistently represented, including *endocytic vesicle lumen* (OR = 14.65), *endocytic vesicle* (OR = 2.33), and *vesicle* (OR = 4.16). Fifth, nitric oxide signaling and proteostasis pathways were also conserved, encompassing *positive regulation of nitric oxide biosynthetic process* (OR = 9.15), *regulation of nitric oxide biosynthetic process* (OR = 6.75), and *ubiquitin protein ligase binding* (OR = 3.00). Together, these results indicate that astrocytes adopt a conserved reactive phenotype across cortical and striatal regions in TD, characterized by neurovascular remodeling, lipid dyshomeostasis, oxidative stress responses, and altered vesicular trafficking.

Oligodendrocytes exhibited enrichment across four principal functional domains. First, intracellular second messenger and calcium signaling was highlighted by *cyclase activator activity* (OR = 35.44), *adenylate cyclase regulator activity* (OR = 23.65), *positive regulation of ryanodine-sensitive calcium-release channel activity* (OR = 17.72), *detection of calcium ion* (OR = 16.91), and *regulation of presynaptic cytosolic calcium levels* (OR = 5.06), suggesting dysregulation of coordinated cAMP and calcium-dependent intracellular signaling in oligodendrocytes across both regions. Second, vesicular trafficking, synaptic structure, and presynaptic signaling were consistently enriched, including *presynaptic endocytosis* (OR = 23.62), *clathrin-sculpted gamma-aminobutyric acid transport vesicle* (OR = 17.72), *extrinsic component of presynaptic membrane* (OR = 14.17), *synaptic signaling via neuropeptide* (OR = 14.17), *synaptic vesicle cycle* (OR = 5.94), *presynaptic active zone cytoplasmic component* (OR = 5.56), *synapse* (OR = 5.45), and *postsynaptic density intracellular component* (OR = 5.21), pointing to conserved alterations in vesicle-mediated GABAergic transmission and synaptic organization; these presynaptic terms may reflect oligodendrocyte-neuron interactions at the synaptic interface rather than direct presynaptic function. Third, mitochondrial and oxidative stress pathways were represented by *regulation of mitochondrial membrane potential* (OR = 4.90), *response to reactive oxygen species* (OR = 4.19), and *oxidative phosphorylation* (OR = 3.65), consistent with metabolic adaptation and mitochondrial stress in oligodendrocytes across both regions. Fourth, proteostasis, biosynthetic, and cytoskeletal processes were also conserved, including *positive regulation of biosynthetic process* (OR = 3.08), *protein stabilization* (OR = 2.49), *ubiquitin protein ligase binding* (OR = 2.47), *axon guidance* (OR = 2.80), *microtubule* (OR = 2.14), and *polymeric cytoskeletal fiber* (OR = 1.83), though these latter terms showed modest enrichment and likely reflect broader structural processes. Downregulated signals in oligodendrocytes yielded only two overlapping GO terms across the two independent datasets: *chromatin remodeling* (OR = 2.06) and *chromatin organization* (OR = 1.71), suggesting a limited but potentially reproducible signal related to chromatin-related processes. Taken together, the transcriptional profile of oligodendrocytes across cortical and striatal regions in TD is characterized by upregulation of activity-associated programs involving second messenger and calcium signaling, vesicle-mediated GABAergic transmission, and mitochondrial metabolic adaptation, accompanied by a limited but potentially reproducible downregulation of chromatin remodeling and organization processes, suggesting enhanced functional engagement coupled with a subtle reduction in transcriptional plasticity.

Pathway enrichment in OPCs revealed five dominant functional domains. First, synaptic organization and interaction pathways showed the strongest enrichment, including *presynaptic endocytosis* (OR = 120.91), *anchored component of presynaptic membrane* (OR = 30.04), *extracellular matrix of synaptic cleft* (OR = 29.86), *anchored component of postsynaptic membrane* (OR = 23.89), *perisynaptic extracellular matrix* (OR = 23.89), *postsynaptic modulation of chemical synaptic transmission* (OR = 7.50), and *vesicle* (OR = 8.31), collectively pointing to conserved alterations in synaptic interface organization and vesicle-mediated signaling; as noted for oligodendrocytes, these synaptic terms likely reflect OPC-neuron interactions rather than direct synaptic function. Second, calcium and calcineurin-dependent signaling was represented by *calcineurin-mediated signaling* (OR = 25.90) and *calcium ion binding* (OR = 4.40), suggesting dysregulation of calcium-dependent intracellular pathways across both regions. Third, metabolic and energetic pathways were consistently enriched, including *regulation of ATP biosynthetic process* (OR = 24.03), *unsaturated fatty acid biosynthetic process* (OR = 18.48), and *ATP metabolic process* (OR = 6.13), consistent with increased energetic and lipid biosynthetic demand in OPCs across cortical and striatal regions. Fourth, immune and stress-response pathways were represented by *cellular response to interferon-beta* (OR = 12.01), *MHC class II protein complex binding* (OR = 10.92), and *cellular response to chemical stress* (OR = 5.05), alongside proteostasis-related processes including *protein stabilization* (OR = 4.84) and *ubiquitin protein ligase binding* (OR = 4.83), suggesting a shared innate immune and stress-adaptive signature. Fifth, developmental and structural pathways were also enriched, including *neuron projection development* (OR = 4.99) and *central nervous system development* (OR = 3.04), though these terms showed modest enrichment and may reflect a broader state of cellular plasticity and responsiveness rather than active developmental remodeling. Together, these results indicate that OPCs exhibit a conserved activity-responsive transcriptional program across cortical and striatal regions in TD, characterized by synaptic interface remodeling, calcium-dependent signaling, metabolic adaptation, and innate immune activation.

Five overarching functional themes emerged from microglia enrichment analysis. First, proteostatic and catabolic processes were highlighted by *modification-dependent macromolecule catabolic process* (OR = 36.92), *cytoplasmic translation* (OR = 22.84), *ribosomal small subunit assembly* (OR = 15.38), *ATP biosynthetic process* (OR = 9.22), and *ubiquitin protein ligase binding* (OR = 5.64), consistent with increased biosynthetic and proteostatic demand in reactive microglia across both regions. Second, structural and synaptic remodeling was represented by *cell junction disassembly* (OR = 30.62), *extrinsic component of presynaptic membrane* (OR = 27.62), and *cadherin binding* (OR = 3.00), suggesting conserved alterations in cell adhesion and synaptic interface organization. Third, immune activation pathways were consistently enriched, including *negative regulation of macrophage differentiation* (OR = 30.62), *humoral immune response mediated by circulating immunoglobulin* (OR = 22.97), *positive regulation of interleukin-4 production* (OR = 9.86), *positive regulation of nitric oxide biosynthetic process* (OR = 14.13), and *positive regulation of cytokine production* (OR = 2.26); the enrichment of *negative regulation of macrophage differentiation* is particularly noteworthy, as it may reflect a shift away from homeostatic microglial identity toward a disease-associated state conserved across both regions. Fourth, lipid and lipoprotein metabolism was represented by *low-density lipoprotein particle* (OR = 23.02), *lipoprotein metabolic process* (OR = 22.97), and *very-low-density lipoprotein particle remodeling* (OR = 22.97), consistent with immunometabolic reprogramming of microglia in TD. Fifth, vesicular and endo-lysosomal trafficking pathways were robustly enriched, including *autolysosome* (OR = 13.12), *endosome lumen* (OR = 8.78), *endocytic vesicle lumen* (OR = 7.67), *transport vesicle membrane* (OR = 4.35), *phagocytic vesicle membrane* (OR = 4.19), and *positive regulation of endocytosis* (OR = 4.16), collectively pointing to enhanced vesicular processing and phagocytic activity. In contrast to the upregulation of vesicular and immune pathways, downregulated signals in microglia were enriched for GTPase regulatory functions (*GTPase regulator activity* OR=3.73, *guanyl nucleotide exchange factor activity*, OR=4.12), indicating reduced regulation of cytoskeletal and trafficking processes. Taken together, the transcriptional profile of microglia across cortical and striatal regions in TD reflects a shift toward a reactive and immunometabolically active state, characterized by upregulation of vesicular processing, immune signaling, and lipid metabolic pathways, coupled with downregulation of GTPase regulatory functions — suggesting enhanced inflammatory and phagocytic activity alongside reduced homeostatic control of cytoskeletal and trafficking dynamics.

In interneurons, enriched pathways were *proton-transporting V-type ATPase, V0 domain* and *vacuolar proton-transporting V-type ATPase, V0 domain* (both OR = 26.03), which likely represent hierarchically related GO terms converging on the same biological process, V-type ATPase–mediated luminal acidification of vesicular compartments. Additional shared enrichment was observed for *cation-transporting ATPase complex* (OR = 3.47), *vesicle* (OR = 1.64), and *microtubule cytoskeleton* (OR = 1.34), although these terms showed modest enrichment and likely reflect broader, non-specific cellular processes. Downregulated signals in interneurons yielded 14 overlapping GO terms across the two independent datasets, converging predominantly on **s**ynaptic structure and function. The most strongly enriched shared terms included *synaptic vesicle cycle* (OR = 51.03), *extrinsic component of postsynaptic membrane* (OR = 34.19), *presynapse assembly* (OR = 25.64), *synapse assembly* (OR = 25.51), and *structural constituent of postsynaptic density* (OR = 18.55), alongside terms related to presynaptic regulation such as *voltage-gated ion channel activity involved in regulation of presynaptic membrane potential* (OR = 13.61), *presynaptic active zone cytoplasmic component* (OR = 10.74), *regulation of synaptic vesicle exocytosis* (OR = 9.04), and *regulation of postsynaptic membrane neurotransmitter receptor levels* (OR = 7.14). Additional terms with more modest enrichment included *integral component of presynaptic membrane* (OR = 6.92), *integral component of postsynaptic density membrane* (OR = 3.76), *postsynaptic density* (OR = 3.30), *axon* (OR = 3.99), and *neuron projection* (OR = 2.93). The convergence of these synaptic terms across independent cohorts with distinct gene sets points to a reproducible signature of synaptic disorganization and impaired inhibitory neurotransmission as a shared feature of interneuron pathology in TD. Taken together, the transcriptional profile of interneurons across cortical and striatal regions in TD is characterized by a dissociation between upregulated vesicle acidification machinery and widespread downregulation of synaptic structural and functional components. This opposing pattern suggests a maladaptive state in which vesicular processing is maintained while the synaptic architecture required for effective inhibitory neurotransmission is progressively dismantled, a signature conserved across independent cohorts and brain regions, pointing to interneuron synaptic disorganization as a reproducible and potentially central feature of TD pathology.

### SUPPLEMENTAL METHODS AND MATERIALS

**Human Brain Collection and Donor Characterization.** Brain tissue samples from the dorsolateral prefrontal cortex (DLPFC; Brodmann area 9, BA9) were obtained from the NIH NeuroBioBank (NBB) Brain Tissue Resource Center (BTR) at Harvard University. Brain samples from individuals with Tourette disorder (TD) were collected through the Tourette Association of America (TAA). Cases with known or suspected neurological diseases or prolonged hypoxia were excluded. The right hemisphere of each brain was coronally sectioned, promptly frozen, and stored at -80°C in accordance with protocols established by the NIH National Brain Bank. The BA9 region was identified based on anatomical landmarks, ensuring the exclusion of white matter. No evidence of pathological atrophy was observed across any brain regions. Consent for tissue donation was obtained from the next of kin, following ethical and legal guidelines. Diagnoses were confirmed by experienced research clinicians through structured interviews with family members or by reviewing prior medical records. Control subjects were screened using the same methodology to ensure the absence of psychiatric disorders.

Postmortem tissue samples from BA9 (30.18 ± 9.55 mg per sample) were analyzed, alongside clinical data including age, sex, brain pH, and postmortem interval (PMI). The final cohort consisted of ten male individuals, including five TD cases and five age-matched neurotypical controls. Unpaired two-sample t tests revealed no significant differences between groups in either age (controls: 25.0 ± 3.5 years; TD: 24.8 ± 5.9 years; $t(8) = 0.14$, $P = 0.89$) or PMI (controls: 20.6 ± 2.4 h; TD: 19.0 ± 4.4 h; $t(8) = 0.32$, $P = 0.76$), indicating that the two groups were well matched for these demographic variables (**Table 1**).

**Tissue Processing and Nuclei Isolation.** Brain specimens were gently thawed in 2 mL of ice-cold phosphate-buffered saline (PBS) without $Ca^{2+}$ and $Mg^{2+}$, supplemented with 0.54 uM Nectostatin (ThermoFisher Scientific, Inc.#J65341), 1 uM HPN-07 (Sigma-Aldrich, SML2163), 0.32 uM sodium hydroxybutyrate (ThermoFisher Scientific, A1161314), 78 nM Q-VD-Oph (Sigma-Aldrich, SML0063), and 0.2 U/mL Ribolock RNase inhibitor (Life Technologies, EO0381). The tissue, maintained on dry ice within a 60-mm Petri dish, was finely minced into fragments (<2 mm) using a sterile razor blade. Fragmented tissue was then transferred into a 2 mL Dounce homogenizer containing 1.5 mL of nuclei extraction buffer (Miltenyi Biotec, 130-128-024) supplemented with 0.2 U/mL Ribolock RNase inhibitor.

The homogenization procedure was conducted as follows: (1) ten strokes with pestle A; (2) ten strokes with pestle B, followed by a five-minute incubation on ice; (3) ten additional strokes with pestle B, followed by another five-minute incubation on ice; and (4) a final set of ten strokes with pestle B. The homogenate was then centrifuged at 600 x g for five minutes, resuspended in 400 µL of PBS containing 2% bovine serum albumin (BSA), and sequentially filtered through a 70-um FLOWMI filter for initial clarification, followed by centrifugation through a 500 µL 1 M sucrose solution, resuspension in 250 µL of PBS containing 2% BSA, and final filtration through a 40-µm FLOWMI filter. The purified nuclei were submitted to the Huntsman Cancer Institute High-Throughput Genomics Core for 10x Genomics library construction and sequencing.

**Single-Nucleus RNA Sequencing Data Processing.** Raw sequencing data were processed using the 10x Genomics Cell Ranger Single Cell pipeline [8] with default settings and alignment to the GRCh38 reference genome. Quality control and downstream analyses were conducted in Seurat (version 4.1.0) using the following filtering criteria: (1) nuclei expressing fewer than 1,000 detected genes or exhibiting more than 5% mitochondrial gene content were removed; (2) mitochondrial and hemoglobin genes were excluded [9–12]. After normalization, variable feature selection, and scaling, all 72,340 nuclei underwent principal component analysis, neighborhood graph construction, SCTransformation-based normalization, and clustering.

**Cell-Type Annotation, Differential Expression, and Functional Profiling.** A custom reference human BA9 annotation dataset and UMAP was generated as follows. Processed data files (matrix.csv and metadata.csv) were downloaded from the PsychENCODE Knowledge Portal located on Synapse [13] and loaded into Seurat. The class, subclass, and cluster labels from the metadata file were retained for cell and cluster annotations. The data were processed with NormalizeData, FindVariableFeatures, and ScaleData. The Azimuth R package (version 0.5) was used for automated cell-type classification [14]. AzimuthReference was used to generate ref.Rds and idx.annoy files, creating a custom human BA9 reference. Single-nuclei data from this study were similarly processed and annotated using RunAzimuth, then projected onto the custom reference UMAP. Results were validated by manual inspection of canonical marker genes.

The annotations of the cell types at the subclass level contained a large "IT" (intra-telencephalic) group; therefore, we combined layer annotations from the cluster-level with the subclass-level annotations to annotate excitatory neuronal subtypes, resolving intra-telencephalic populations into layer-specific clusters.

Differentially expressed genes (DEGs) were identified using the FindAllMarkers function on SCT-normalized data with false discovery rate (FDR) < 0.05 and a minimum 1.5-fold expression change over background. Gene Ontology (GO) enrichment analyses were conducted using enrichR (version 3.4) [15–17], with top-ranked pathways selected based on adjusted P values. For heatmap generation, DEGs with a P value < 0.0125 were selected for GO analysis; the resulting GO terms were ranked by adjusted P values in ascending order, and the top 20 unique pathways were identified and their expression patterns across all clusters were analyzed.

**Stress-Responsive and Immediate Early Gene Analyses.** To assess stress-related transcriptional engagement, DEGs were compared with dexamethasone-responsive gene sets derived from iPSC neurons [1]. Module scores were computed for each cell type. Immediate early gene (IEG) enrichment was evaluated using a curated reference set [7], with statistical significance assessed by Fisher's exact test. Potential confounding effects of postmortem interval were examined using Spearman correlations between PMI and IEG expression at single-nucleus, donor, and cell-type levels, with FDR correction applied.

**Cross-Disorder Transcriptional Comparisons.** To contextualize TD-associated transcriptional changes within a broader psychiatric framework, DEGs were compared with published single-nucleus datasets from major depressive disorder (MDD) and posttraumatic stress disorder (PTSD) affecting the DLPFC [1]. Gene overlap was quantified using the R package GeneOverlap (version 1.42.0) [18], stratified by cell type and direction of expression change. For each triplet combination of TD, MDD, and PTSD cell types, gene sets were extracted from differential expression results, stratified by increased or decreased expression. The intersection of gene sets across the three conditions was computed for both positively and negatively regulated genes, and overlaps were visualized using Venn diagrams.

**Integration with TD Genetic Architecture.** To determine whether transcriptional alterations intersect with genetic risk, BA9 DEGs were compared with gene sets derived from the TD GWAS meta-analysis [2]. Analyses included: (i) genes harboring eQTLs and sQTLs linked to TD risk variants; (ii) gene-level associations from MAGMA [19], including standard (c-MAGMA), Hi-C chromatin interaction-informed (h-MAGMA), and eQTL-informed (e-MAGMA) models; and (iii) cell-type-specific associations from brainSMR analyses [20]. Overlap between GWAS-derived gene sets and DEGs was assessed using Fisher's exact tests across all cell types and expression directions, with the background defined as all detected genes.

**Cross-Regional Transcriptional Comparisons.** To assess cross-regional convergence across CSTC circuitry, BA9 DEGs were compared with published striatal snRNA-seq data from TD [3] using the GeneOverlap framework [18], applied across all pairwise cell-type comparisons separately for upregulated and downregulated genes. Because certain cell types are region-specific (striatal projection neurons in basal ganglia and excitatory neurons in cortex), the most informative comparisons were limited to cell classes represented in both regions. Shared gene sets were analyzed for pathway enrichment using enrichR.

**Pseudotime and Maturation Trajectory Analysis.** Pseudotime analysis was performed using Monocle3 (version 1.4.26) [21, 22] to investigate the maturation trajectories of specific neuronal populations. After clustering neuronal populations, root cells were selected based on canonical marker gene expression to initiate the trajectory. The Monocle3 algorithm inferred the progression of cell states along pseudotime, assigning each cell a pseudotime value reflecting its position in the maturation trajectory. Pseudotime trajectories were compared between TD and control groups for each neuronal subtype using the Wilcoxon rank-sum test. A rightward shift in pseudotime indicated accelerated maturation, while a leftward shift suggested delayed maturation. DEGs along pseudotime trajectories were identified by comparing gene expression profiles across maturation stages and subjected to GO enrichment analysis.

**Statistical Analysis.** All statistical analyses were conducted using R (version 4.1.0), with the exception of BA9 expression specificity analyses, which were conducted in Python 3. Differential gene expression between TD and control groups was assessed using the Wilcoxon rank-sum test, with genes exhibiting a log2 fold-change > 1.5 and a false discovery rate (FDR) < 0.05 considered statistically significant. Pseudotime comparisons were made using the Wilcoxon rank-sum test. For GO analysis, a P value < 0.05 was used to select differentially expressed genes, and the top 20 enriched pathways were visualized as heatmaps. GWAS convergence analyses used Fisher's exact tests (one-sided) and Mann-Whitney U tests as described above. All visualizations, including UMAP plots, pseudotime trajectories, and heatmaps, were generated using Seurat and ggplot2. Results are presented as mean ± SEM, and statistical significance was set at P < 0.05 for all analyses unless otherwise specified.

**SUPPLEMENTARY TABLES**

| | Astro | PN | IN | Microglia | Oligo | OPC |
|---|---|---|---|---|---|---|
| Astro | 36.41 | 1.30 | 1.01 | 0.91 | 0.00 | 1.82 |
| PN | 0.00 | 100.00 | 0.21 | 0.00 | 0.00 | 0.00 |
| IN | 0.00 | 0.38 | 30.54 | 0.00 | 0.00 | 0.51 |
| Microglia | 0.00 | 0.00 | 0.00 | 100.00 | 0.00 | 0.00 |
| Oligo | 0.54 | 0.00 | 0.00 | 0.00 | 100.00 | 3.25 |
| OPC | 0.00 | 0.00 | 3.64 | 0.00 | 0.00 | 38.22 |

**Supplementary Table 1:** Cross-dataset correspondence of cell-type cluster assignments with [1]. Odds ratios quantifying the correspondence between cell-type clusters identified in the present dataset (rows) and reference dataset annotations (columns). Higher odds ratios indicate stronger correspondence between clusters. Cell shading reflects the magnitude of the odds ratio, ranging from yellow (higher values) to blue (lower values). Cells outlined in red denote statistically significant associations after correction for multiple comparisons ($P < 0.001$). Abbreviations: Astro, astrocytes; PN, pyramidal neurons; IN, interneurons; Oligo, oligodendrocytes; OPC, oligodendrocyte precursor cells.

| Clusters | Overlap | P value | Odds ratio |
|---|---|---|---|
| Global | 53 | 7.5845E-09 | 2.913379105 |
| Pyramidal neurons | 45 | 7.475E-09 | 3.114566705 |
| Interneurons | 28 | 0.026884042 | 1.564550114 |
| Astrocytes | 23 | 1.39909E-13 | 8.693377246 |
| Oligodendrocytes | 15 | 0.00011658 | 3.362769287 |
| Oligodendrocyte precursor cells | 13 | 2.61945E-11 | 15.32806613 |
| Microglia | 31 | 8.65096E-25 | 16.38797939 |

**Supplementary Table 2. Widespread enrichment of upregulated IEGs in DLPFC cell types in TD.** All major dorsolateral prefrontal cortex (DLPFC) cell populations exhibited significant enrichment of upregulated immediate early genes (IEGs, curated list obtained by [7]) in individuals with Tourette disorder (TD) compared with neurotypical controls, indicating broad activation of stress-responsive and transcriptionally induced pathways across both neuronal and glial lineages. The table summarizes the enrichment statistics for each cell class, including the number of overlapping upregulated IEGs (*Overlap*), the statistical significance of enrichment (*P* value, Fisher's exact test), and the *Odds ratio* representing the magnitude of enrichment.

### SUPPLEMENTARY DATA

Data S1: Data from single-nucleus transcriptomic analyses of the dorsolateral prefrontal cortex, showing the genes upregulated in TD patients compared with neurotypical individuals across all identified cell clusters, along with the corresponding GO enrichment results.

Data S2: Data from single-nucleus transcriptomic analyses of the dorsolateral prefrontal cortex, showing the genes downregulated in TD patients compared with neurotypical individuals across all identified cell clusters, along with the corresponding GO enrichment results.

Data S3: Data from single-nucleus transcriptomic analyses of the dorsolateral prefrontal cortex, showing the differentially expressed genes (DEGs) in the comparison of TD patients and neurotypical individuals across all identified cell clusters, along with the corresponding GO enrichment results.

Data S4: Upregulated genes in TD patients compared with neurotypical individuals across all identified excitatory neuron clusters, along with the corresponding GO enrichment results.

Data S5: Downregulated genes in TD patients compared with neurotypical individuals across all identified excitatory neuron clusters, along with the corresponding GO enrichment results.

Data S6: DEGs in the comparison of TD patients and neurotypical individuals across all identified excitatory neuron clusters, along with the corresponding GO enrichment results.

Data S7: Wilcoxon rank-sum test results for group comparisons of pseudotime values in excitatory neurons.

Data S8: Upregulated genes in TD patients compared with neurotypical individuals across all identified inhibitory neuron clusters, along with the corresponding GO enrichment results.

Data S9: Downregulated genes in TD in TD patients compared with neurotypical individuals across all identified inhibitory neuron clusters, along with the corresponding GO enrichment results.

Data S10: DEGs in the comparison of TD patients and neurotypical individuals across all identified inhibitory neuron clusters, along with the corresponding GO enrichment results.

Data S11: Wilcoxon rank-sum test results for group comparisons of pseudotime values in interneurons.

Data S12: Expression profiles of dexamethasone-responsive genes, derived from [1], showing their differential expression between TD patients and neurotypical individuals across all identified clusters.

Data S13: List of positively and negatively overlapping differentially expressed genes (DEGs) shared between MDD and PTSD (derived from [1]) and TD patient-control comparisons, across all identified cell-type clusters.

Data S14: Expression profiles of immediate-early genes, derived from [7], showing their differential expression between TD patients and neurotypical individuals across all identified clusters.

DataS15: Overlap with Strom [2] datasets (sQTL, eQTL, c-MAGMA, h-MAGMA, e-MAGMA).

DataS16: Overlap with Strom [2] datasets (brainSMR).

Data S17: Comparison between upregulated gene in TD patients compared with data derived from [3] across all identified clusters.

Data S18: Comparison between upregulated GO terms in TD patients compared with data derived from [3] across all identified clusters.

Data S19: Comparison between downregulated gene in TD patients compared with data derived from [3] across all identified clusters.

Data S20: Comparison between downregulated GO terms in TD patients compared with data derived from [3] across all identified clusters.

### SUPPLEMENTARY REFERENCES


1. Chatzinakos C, Pernia CD, Morrison FG, Iatrou A, McCullough KM, Schuler H, et al. Single-Nucleus Transcriptome Profiling of Dorsolateral Prefrontal Cortex: Mechanistic Roles for Neuronal Gene Expression, Including the 17q21.31 Locus, in PTSD Stress Response. Am J Psychiatry. 2023;180:739–754.

2. Strom NI, Halvorsen MW, Grove J, Ásbjörnsdóttir B, Luðvígsson P, Thorarensen Ó, et al. Genome-Wide Association Study Meta-Analysis of 9619 Cases With Tic Disorders. Biol Psychiatry. 2025;97:743–752.

3. Wang Y, Fasching L, Wu F, Suvakov M, Huttner A, Berretta S, et al. Interneuron Loss and Microglia Activation by Transcriptome Analyses in the Basal Ganglia of Tourette Disorder. Biological Psychiatry. 2025;98:260–270.

4. Kalanithi PSA, Zheng W, Kataoka Y, DiFiglia M, Grantz H, Saper CB, et al. Altered parvalbumin-positive neuron distribution in basal ganglia of individuals with Tourette syndrome. Proc Natl Acad Sci U S A. 2005;102:13307–13312.

5. Kataoka Y, Kalanithi PSA, Grantz H, Schwartz ML, Saper C, Leckman JF, et al. Decreased number of parvalbumin and cholinergic interneurons in the striatum of individuals with Tourette syndrome. J Comp Neurol. 2010;518:277–291.

6. Haber SN, Liu H, Seidlitz J, Bullmore E. Prefrontal connectomics: from anatomy to human imaging. Neuropsychopharmacol. 2022;47:20–40.

7. Wu YE, Pan L, Zuo Y, Li X, Hong W. Detecting Activated Cell Populations Using Single-Cell RNA-Seq. Neuron. 2017;96:313-329.e6.

8. Zheng GXY, Terry JM, Belgrader P, Ryvkin P, Bent ZW, Wilson R, et al. Massively parallel digital transcriptional profiling of single cells. Nat Commun. 2017;8:14049.

9. Butler A, Hoffman P, Smibert P, Papalexi E, Satija R. Integrating single-cell transcriptomic data across different conditions, technologies, and species. Nat Biotechnol. 2018;36:411–420.

10. Romanov RA, Tretiakov EO, Kastriti ME, Zupancic M, Häring M, Korchynska S, et al. Molecular design of hypothalamus development. Nature. 2020;582:246–252.

11. Satija R, Farrell JA, Gennert D, Schier AF, Regev A. Spatial reconstruction of single-cell gene expression data. Nat Biotechnol. 2015;33:495–502.

12. Stuart T, Butler A, Hoffman P, Hafemeister C, Papalexi E, Mauck WM, et al. Comprehensive Integration of Single-Cell Data. Cell. 2019;177:1888-1902.e21.

13. Zintel T. Cellular Diversity in Human Subgenual Anterior Cingulate and Dorsolateral Prefrontal Cortex by Single-Nucleus RNA-sequencing. 2022.

14. Hao Y, Hao S, Andersen-Nissen E, Mauck WM, Zheng S, Butler A, et al. Integrated analysis of multimodal single-cell data. Cell. 2021;184:3573-3587.e29.

15. Chen EY, Tan CM, Kou Y, Duan Q, Wang Z, Meirelles GV, et al. Enrichr: interactive and collaborative HTML5 gene list enrichment analysis tool. BMC Bioinformatics. 2013;14:128.

16. Kuleshov MV, Jones MR, Rouillard AD, Fernandez NF, Duan Q, Wang Z, et al. Enrichr: a comprehensive gene set enrichment analysis web server 2016 update. Nucleic Acids Res. 2016;44:W90–W97.

17. Xie Z, Bailey A, Kuleshov MV, Clarke DJB, Evangelista JE, Jenkins SL, et al. Gene Set Knowledge Discovery with Enrichr. Current Protocols. 2021;1:e90.

18. Li Shen MS <Shenli SC. GeneOverlap. 2017.

19. Leeuw CA de, Mooij JM, Heskes T, Posthuma D. MAGMA: Generalized Gene-Set Analysis of GWAS Data. PLOS Computational Biology. 2015;11:e1004219.

20. Zhu Z, Zhang F, Hu H, Bakshi A, Robinson MR, Powell JE, et al. Integration of summary data from GWAS and eQTL studies predicts complex trait gene targets. Nat Genet. 2016;48:481–487.

21. Cao J, Spielmann M, Qiu X, Huang X, Ibrahim DM, Hill AJ, et al. The single-cell transcriptional landscape of mammalian organogenesis. Nature. 2019;566:496–502.

22. Trapnell C, Cacchiarelli D, Grimsby J, Pokharel P, Li S, Morse M, et al. The dynamics and regulators of cell fate decisions are revealed by pseudotemporal ordering of single cells. Nat Biotechnol. 2014;32:381–386.
